# Decoding Cell-level Circadian Phase and Rhythm-state Remodeling in Single-cell Transcriptomes With scRhythmBayes

**DOI:** 10.64898/2026.10.18.759931

**Authors:** Qian Zhang, Chenyu Li, Ioannis Konstantinos Deligiannis, Wenhong Jiao, Weigang Liu, Celia Pilar Martinez-Jimenez, Zhibin Wang, Kenneth A. Dyar, Liwei Zhang, Pin Sun

**Affiliations:** Affiliated Hospital of Qingdao University, Cardiac Ultrasound Department, Qingdao, China; Institute of Biotechnology and Biomedicine (BIOTECMED), Department of Biochemistry and Molecular Biology, University of Valencia, Burjassot, Spain; Metabolic Physiology, Institute for Diabetes and Cancer, Helmholtz Munich, German Research Center for Environmental Health and German Center for Diabetes Research (DZD), Munich, Germany

**Keywords:** circadian phase inference, rhythmic gene expression, differential rhythmicity, dynamic time warping, cell-type-specific rhythmic programs

## Abstract

Current circadian omics analysis tools have been developed primarily for bulk time-series. However, heterogeneous single-cell datasets require analysis and interpretation of rhythmic programs across different cells, cell types, and biological conditions. In these datasets, cellular circadian phase is usually unobserved, and sample-level zeitgeber time provides temporal information but does not define the exact phase of every cell. Existing methods address only selected aspects of this problem and do not model how cell-level phase evidence informs gene-level rhythmic parameters and rhythm-state remodeling. Here we present scRhythmBayes, a new probabilistic algorithm for phase-resolved circadian analysis in heterogeneous single-cell transcriptomes. scRhythmBayes combines a seed-gene count likelihood with a stratum-aware circular prior to infer cell-level circadian phase, incorporates phase support into gene-level rhythmic modeling, and assigns posterior probabilities to interpretable rhythm-state categories, including rhythm gain, rhythm loss, amplitude remodeling and phase shifts. In heterogeneous simulations, scRhythmBayes achieved higher phase-recovery accuracy than existing phase-inference methods and maintained balanced performance across rhythmic gene detection, parameter recovery, and rhythm-state classification. Stress tests defined practical requirements for seed-gene support, zeitgeber anchoring, temporal sampling density, and cell number. In mouse liver, scRhythmBayes revealed distinct circadian phase occupancy and rhythm-state organization between periportal and pericentral hepatocytes, supporting a temporal dimension of liver zonation. It also identified cell-type-specific rhythmic markers that were largely distinct from conventional static markers, indicating that cellular identity includes temporal programs not captured by average expression. In human heart disease data, phase-coordination and phase-distribution analyses identified cell-type-selective temporal disorganization rather than uniform circadian disruption across the tissue. scRhythmBayes therefore provides a statistically grounded approach for resolving circadian organization across cells, genes, cell types and biological conditions.

**Highlights:**

- scRhythmBayes establishes circadian phase as a latent cellular coordinate for single-cell transcriptomic analysis.
- scRhythmBayes introduces original probabilistic models for cell-phase inference, phase-confidence-aware gene modeling, and rhythm-state classification.
- Cell-type-specific rhythmic programs reveal a temporal dimension of cellular identity inaccessible to static analyses.
- Phase-resolved single-cell analyses identify circadian remodeling across tissues, physiological states, and disease.

## Introduction

Circadian rhythms coordinate transcriptional, metabolic and physiological processes across the 24-h day, and rhythmic gene expression has been reported across many mammalian and human tissues^1,2^. Transcriptome-scale circadian analysis, however, has largely been developed in the context of bulk time-series experiments, where each sample has an observed collection time and gene expression represents an average signal across many cells. In this setting, the primary inferential task is to test whether a gene oscillates over known sampling times, or to compare rhythmic parameters between conditions. Single-cell transcriptomics create a fundamentally different inference problem. Each cell is captured as a sparse and noisy transcriptomic snapshot, its intrinsic circadian phase is usually unobserved, and cells collected at the same zeitgeber time can differ in cell type, biological state, sequencing depth and temporal alignment^3-5^. Thus, single-cell circadian analysis is not simply bulk rhythmicity testing applied to smaller units. It requires dedicated statistical models that infer latent cellular phase and gene-level rhythmic parameters from sparse heterogeneous observations while preserving the circular geometry of circadian time.

Several methods address important components of circadian transcriptome analysis. Tempo performs Bayesian circadian phase inference in single-cell transcriptomes^5^. CYCLOPS reconstructs cyclic order from unordered transcriptomic samples^6^, and Cyclum infers latent periodic processes from single-cell RNA-seq data^4^. However, these methods do not provide a single-cell-specific statistical formulation in which cell-level phase evidence directly informs gene-level rhythmic modeling and differential rhythm-state inference. At the gene level, JTK_CYCLE tests rhythmicity^7^, harmonic regression and related sinusoidal models estimate waveform features such as mesor, amplitude and phase^8^, and dryR supports differential rhythmicity analysis across experimental conditions^9^. These methods have been powerful for bulk or externally timed transcriptomic studies, but they are not designed to infer cell-specific circadian phase from heterogeneous single-cell snapshots. As a result, current approaches do not explicitly model how uncertainty in latent cellular phase should influence downstream estimation of rhythmic parameters and rhythm-state changes. What remains lacking is a single-cell-native probabilistic algorithm that estimates latent cellular phase, incorporates phase confidence into gene-level count modeling, and assigns differential rhythm states from condition-specific rhythmic parameters.

This gap limits biological interpretation in single-cell circadian studies. Most downstream questions are not limited to ordering cells along a cycle. Which cells occupy different regions of circadian phase space? Which genes gain or lose rhythmicity in a given cell type or condition? Which programs change amplitude, shift phase or become temporally disorganized? Which cell populations show disease-associated disruption of circadian coordination? Answering these questions requires dedicated models for cell-level phase inference, phase-confidence-aware gene modeling, rhythm-state classification, and cell-population prioritization.

Here we present scRhythmBayes, a new probabilistic algorithm for phase-resolved circadian analysis in heterogeneous single-cell transcriptomes. scRhythmBayes infers latent circadian phase using a seed-gene count likelihood and a stratum-aware circular mixture prior anchored by sample-level zeitgeber time. It incorporates inferred phase and cell-level phase support into gene-level count models to estimate mesor, amplitude, phase, and rhythmicity, and then assigns posterior support to interpretable rhythm-state categories. To support discovery in multi-cell-type tissues, we further introduce SRB_OVR, a one-versus-rest extension that identifies cell-type-specific rhythmic markers by comparing a target cell type with all remaining cell types pooled as Rest. For multi-condition datasets, scRhythmBayes also provides phase-level prioritization using sample-level phase coordination and dynamic time warping-based phase-distribution alignment.

We benchmarked scRhythmBayes in heterogeneous simulated single-cell data and evaluated its robustness to seed-gene support, zeitgeber anchoring, temporal sampling density and cell number. We then applied scRhythmBayes to mouse liver and human heart disease datasets. In liver, scRhythmBayes resolved rhythm-state remodeling between periportal and pericentral hepatocytes, and identified cell-type-specific rhythmic programs across major liver cell populations. In human heart disease data, phase-coordination and phase-distribution analyses prioritized cardiac cell types with condition-associated temporal disorganization. Together, these analyses establish scRhythmBayes as a new probabilistic algorithm for resolving latent cellular time and interpretable circadian remodeling across cells, genes, cell types, and biological conditions.

## Results

### scRhythmBayes models cell-level circadian phase and rhythm-state remodeling in single-cell transcriptomes

We first outline the scRhythmBayes algorithm and its main inference objectives (Fig. 1). scRhythmBayes is designed for sparse single-cell RNA-seq count matrices with sample-level zeitgeber time annotations and cell-level metadata such as cell type, stratum or condition. The algorithm contains three statistically coupled inference components. The first component infers a posterior circadian phase for each cell using seed-gene counts and sample-level zeitgeber time, together with a cell-level phase-confidence summary. The second component incorporates inferred phase and phase confidence into gene-level count models to estimate mesor, amplitude, phase, and rhythmicity within each condition or stratum. The third component uses condition-specific rhythmic parameters to assign posterior support across eight differential rhythm states, including non-rhythmic, stable rhythm, rhythm gain, rhythm loss, amplitude remodeling, and phase shifts.

**Fig. 1.**
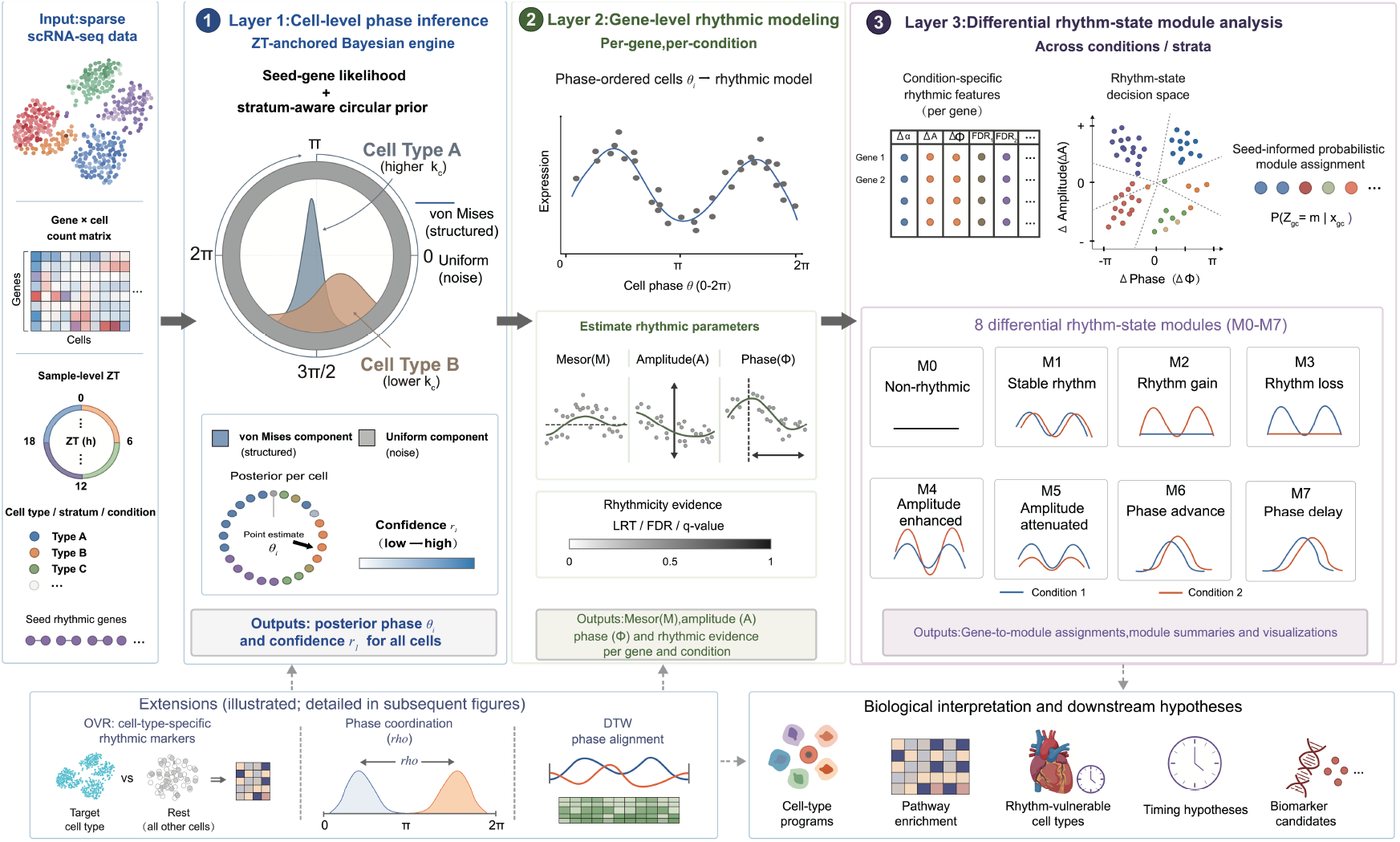
Overview of the scRhythmBayes algorithm for phase-resolved single-cell circadian analysis. scRhythmBayes takes as input a sparse single-cell RNA-seq count matrix, sample-level Zeitgeber time, cell-type or stratum annotations and a panel of seed rhythmic genes. In the first component, cell-level circadian phase is inferred using a seed-gene likelihood together with a stratum-aware circular prior anchored by sample-level Zeitgeber time, producing posterior phase estimates and confidence summaries for individual cells. In the second component, inferred phase and cell-level phase support are incorporated into gene-level count models within each condition or cell population, yielding mesor, amplitude, phase, and rhythmicity evidence. In the third component, condition-specific rhythmic parameters are used to assign posterior support to interpretable differential rhythm states, including non-rhythmic, stable rhythm, rhythm gain and loss, amplitude remodeling, and phase-shift states. Additional extensions support one-versus-rest discovery of cell-type-specific rhythmic programs, sample-level phase coordination based on the mean resultant length ρ and dynamic time warping-based phase-distribution alignment for prioritizing rhythm-vulnerable cell types. These outputs support downstream pathway analysis and biological interpretation of circadian regulation across cell types and conditions.

The inferred phases and gene-level parameters also support two additional analyses designed for heterogeneous tissues and multi-condition datasets. One-versus-rest rhythmic marker analysis identifies cell-type-specific rhythmic programs by comparing a target cell type with the pooled remainder. Phase-distribution summaries, including mean resultant length and dynamic time warping, quantify phase coordination and phase-profile divergence across samples, cell types and conditions. Together, these analyses enable identification of temporally specialized cell populations, cell-type-specific rhythmic programs, and condition-associated temporal disorganization.

### scRhythmBayes accurately recovers cell-level circadian phase in heterogeneous single-cell simulations

We first evaluated whether scRhythmBayes could recover cell-level circadian phase in heterogeneous single-cell data. We simulated four cell types sampled at six symbolic zeitgeber time points (ZT0, ZT4, ZT8, ZT12, ZT16 and ZT20), while introducing continuous within-time-point temporal jitter so that each cell carried a continuous ground-truth circadian time. The simulations incorporated cell-type-specific baseline expression, marker-gene enrichment, modest cell-type-specific phase shifts, library-size variation, Poisson sampling and expression-dependent dropout, producing a sparse and asynchronously sampled benchmark for phase inference.

We compared scRhythmBayes with Tempo, CYCLOPS and Cyclum, which represent Bayesian single-cell circadian phase inference, cyclic transcriptomic ordering and latent periodic process inference, respectively^4-6^. Across the four simulated cell types, scRhythmBayes recovered predicted time distributions that most closely matched the ground truth across ZT bins, whereas the other methods showed stronger distributional distortion, including compression, broadening or systematic displacement of predicted time (Fig. 2a). Circular summaries similarly showed that scRhythmBayes preserved the periodic organization of the simulated 24-h phase structure more faithfully than the competing methods (Fig. 2b).

**Fig. 2.**
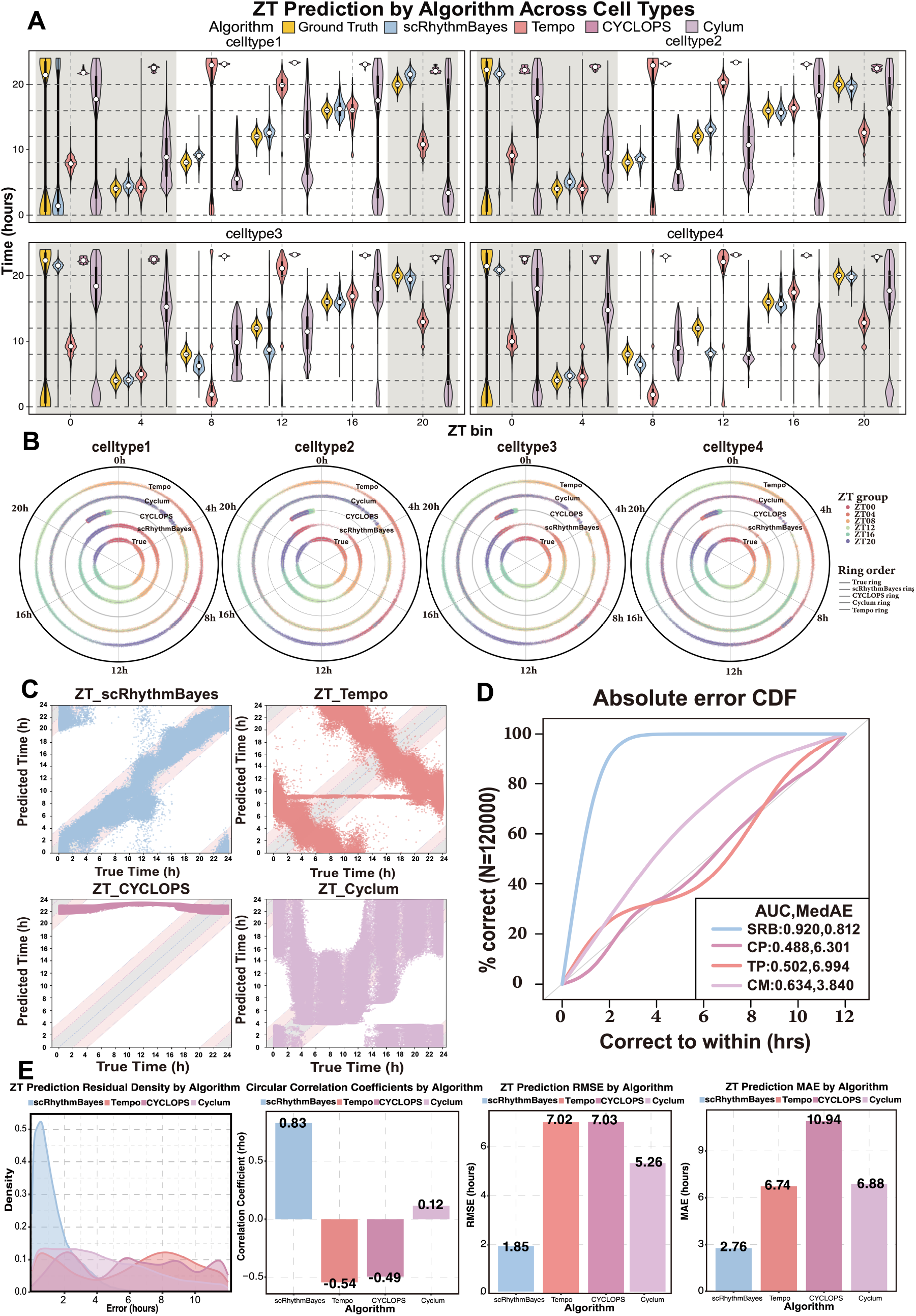
Benchmarking of cell-level circadian phase prediction algorithms in heterogeneous simulated cell types. **a**, Predicted circadian time across six sampling bins (ZT0, ZT4, ZT8, ZT12, ZT16 and ZT20) in four simulated cell types for ground truth, scRhythmBayes, Tempo, CYCLOPS and Cyclum. Violin plots show the distribution of predicted time on the 24-h scale; alternating background shading denotes day and night intervals. **b**, Circular summaries of predicted circadian phase across simulated cell types for Ground Truth and each method. **c**, Scatter plots of predicted versus true circadian time for scRhythmBayes, Tempo, CYCLOPS, and Cyclum. **d**, Empirical cumulative distribution functions (ECDFs) of circular absolute error, with insets showing normalized area under the curve (AUC) and median absolute error (MedAE). **e**, Residual density distributions, circular correlation coefficients, root mean squared error (RMSE), and mean absolute error (MAE) for each method.

At single-cell resolution, predicted-versus-true time plots showed that scRhythmBayes predictions were concentrated near the identity line, whereas Tempo, CYCLOPS and Cyclum showed larger structural deviations from the ground truth (Fig. 2c)^4-6^. Using the empirical cumulative distribution function of circular absolute error, scRhythmBayes achieved the highest normalized AUC (0.920) and the lowest median absolute error (MedAE, 0.812 h). By contrast, Tempo, CYCLOPS and Cyclum achieved AUC values of 0.502, 0.488 and 0.634, with MedAE values of 6.904 h, 6.301 h and 3.840 h, respectively (Fig. 2d)^4-6^. Additional metrics supported the same conclusion: scRhythmBayes showed the strongest circular correlation with the ground truth (ρ = 0.83), and the lowest RMSE (1.85 h) and MAE (2.76 h), compared with substantially higher errors for the other methods (Fig. 2e).

Cell-type-resolved analyses confirmed that the pooled performance was not driven by a single simulated population. Ground-truth time distributions remained well structured within each cell type, and scRhythmBayes retained the most favorable error profiles and predicted-versus-true time structure across cell types (Supplementary Fig. 1a,b,e). Simulated trajectories of clock and clock-controlled genes confirmed that the benchmark contained structured rhythmic signals rather than trivial sampling labels (Supplementary Fig. 1c), and circular phase summaries and cell-type-wise accuracy metrics were consistent with the pooled ranking (Supplementary Fig. 1d,f). These results show that scRhythmBayes recovers latent cell-level circadian phase robustly in the presence of cell-type heterogeneity, temporal jitter and single-cell sparsity.

### scRhythmBayes estimates gene-level rhythmic parameters and differential rhythm states from inferred cellular phase

We next evaluated whether the gene-level model could recover rhythmic parameters and differential rhythm states from inferred cellular phase. Using simulated two-condition single-cell data with known rhythmic truth, we evaluated rhythmic gene detection, recovery of rhythmic parameters, and assignment of genes to differential rhythm-state modules. scRhythmBayes identified 3,497 rhythmic genes and 503 non-rhythmic genes, closely matching the simulation truth of 3,500 rhythmic and 500 non-rhythmic genes (Fig. 3a). JTK underestimated rhythmic gene number, whereas harmonic regression and dryR produced totals closer to the truth but with different error structures^7-9^. Overlap analysis showed a largely shared core of rhythmic genes across methods, together with smaller method-specific call sets (Fig. 3b).

**Fig. 3.**
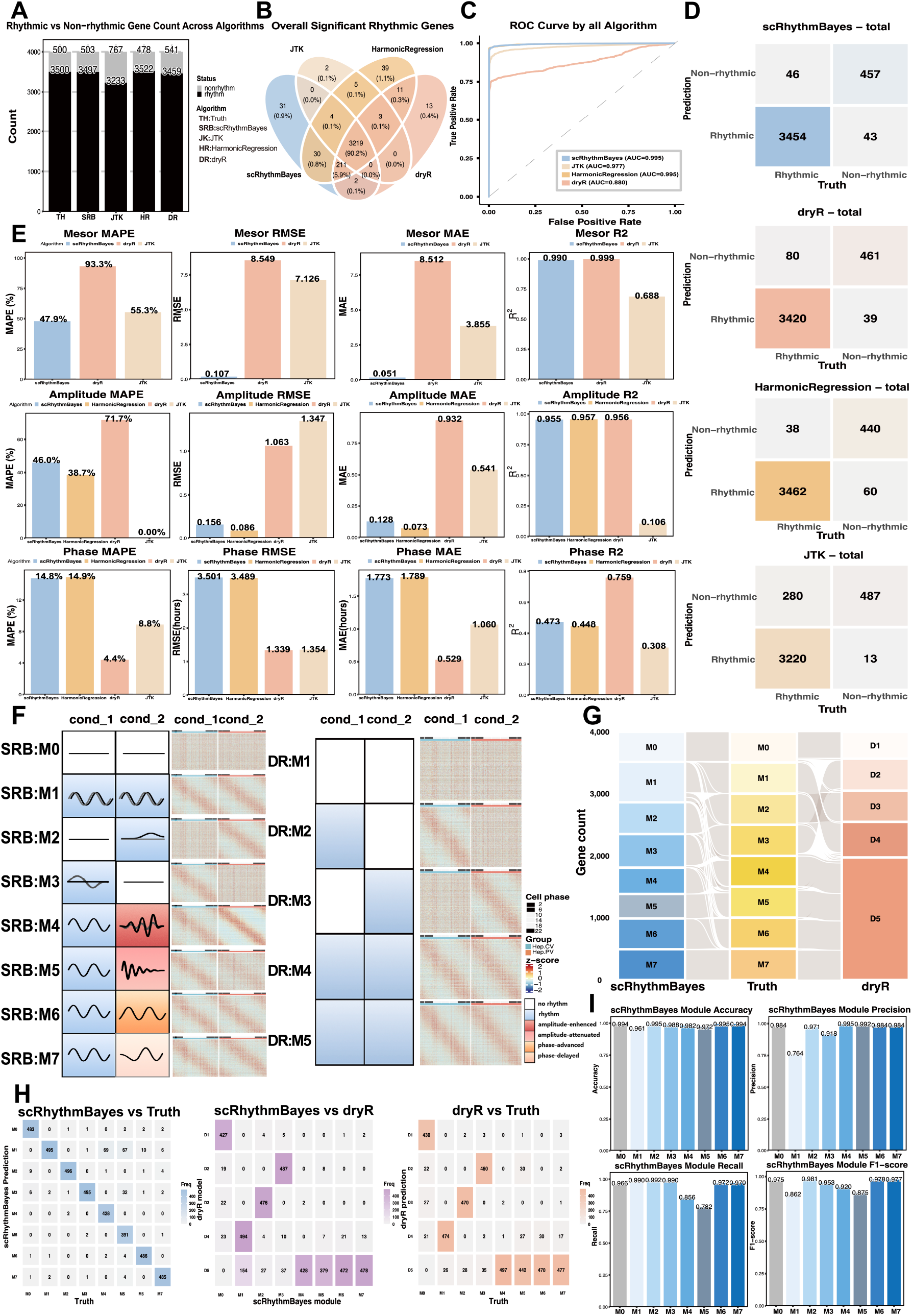
Benchmarking of gene-level rhythmic modeling and module classification in simulated single-cell data. **a**, Bar plots showing the numbers of rhythmic and non-rhythmic genes identified by Ground Truth, scRhythmBayes (SRB), JTK, harmonic regression (HR), and dryR (DR). **b**, Venn diagram summarizing the overlap of significant rhythmic genes identified by SRB, JTK, HR, and DR. **c**, Receiver operating characteristic (ROC) curves for overall rhythmic gene detection across algorithms, with area under the curve (AUC) indicated in the legend. **d**, Confusion matrices comparing predicted and true rhythmic versus non-rhythmic gene labels for SRB, DR, HR, and JTK. **e**, Comparisons of mesor, amplitude, and phase recovery across algorithms, including MAPE, RMSE, MAE, and R^2^. **f**, Representative dynamic patterns and paired heatmaps for SRB modules (M0–M7) and DR models (D1–D5) across the two simulated conditions. **g**, Sankey diagram showing correspondence among SRB module assignments, simulation truth, and DR model labels. **h**, Confusion-matrix summaries comparing SRB predictions with simulation truth, SRB predictions with DR labels, and DR labels with simulation truth. **i**, Module-wise performance summaries for SRB, including accuracy, precision, recall, and F1-score across the eight predefined rhythm-state modules.

ROC analysis showed that scRhythmBayes achieved strong rhythmic gene detection performance (AUC = 0.995), matching harmonic regression and exceeding JTK and dryR (Fig. 3c)^7-9^. Confusion matrices further showed that scRhythmBayes balanced high true-positive recovery with limited false-positive calls, whereas JTK was more conservative and harmonic regression was more permissive (Fig. 3d)^7,8^. These patterns were consistent across the two simulated conditions, where scRhythmBayes retained rhythmic and non-rhythmic gene counts close to the truth and showed stable condition-wise detection performance (Supplementary Fig. 2a–e).

We then evaluated recovery of mesor, amplitude and phase. scRhythmBayes showed the strongest mesor recovery among the methods that provide comparable mesor estimates, and strong amplitude recovery, with low RMSE and MAE and high R^2^ (Fig. 3e). Phase recovery was broadly comparable between scRhythmBayes and harmonic regression, whereas dryR performed well for some phase-related summaries^8,9^. Thus, scRhythmBayes did not dominate every isolated parameter metric, but provided balanced performance across rhythmic gene detection, mesor recovery, amplitude recovery and downstream interpretability. Condition-wise analyses supported the same conclusion across both simulated conditions (Supplementary Fig. 2f).

Finally, we evaluated differential rhythm-state assignment. scRhythmBayes assigns posterior support across eight predefined differential rhythm states: non-rhythmic, stable rhythm, rhythm gain, rhythm loss, amplitude-enhanced, amplitude-attenuated, phase-advanced and phase-delayed states. Each gene is assigned one primary rhythm-state label according to the maximum posterior probability. To retain uncertainty information, scRhythmBayes also reports posterior probabilities for all states and an assignment margin measuring the separation between the top-supported states. Genes with both amplitude and phase differences are therefore represented by their dominant remodeling pattern, while mixed or borderline cases can still be identified from posterior support and margin values. This design provides a compact and interpretable state space for genome-wide differential circadian remodeling. Sankey and confusion-matrix summaries showed that most genes were assigned to the correct simulated module, with misclassifications concentrated between biologically adjacent states (Fig. 3g,h). Module-wise accuracy, precision, recall and F1 score were high for most rhythm-state modules, with lower performance mainly in amplitude-attenuated states where rhythm-state boundaries are intrinsically more continuous (Fig. 3i). Together, these results show that scRhythmBayes uses inferred cellular phase and phase support to estimate gene-level rhythmic parameters and probabilistically resolve differential circadian remodeling.

### scRhythmBayes remains stable across seed-gene, zeitgeber-prior, temporal-sampling and cell-number perturbations

We next tested the robustness of scRhythmBayes to key modeling and experimental factors. Reducing the seed-gene panel from 27 to 20, 15 and 10 genes caused only gradual loss of phase accuracy, whereas the five-gene setting showed a larger decline but retained a visible relationship between predicted and true circadian time (Supplementary Fig. 3a). In contrast, removing the sample-level zeitgeber prior markedly impaired phase recovery: MedAE increased from 0.873 h to 5.962 h and normalized AUC decreased from 0.914 to 0.501 (Supplementary Fig. 3b). These results indicate that zeitgeber time functions as an important circular orientation anchor rather than as a direct observed cell-level phase label.

We then examined the effect of temporal sampling density. Cell-phase inference remained broadly stable when input zeitgeber time points were reduced from 24 to 12, 8, 6 and 4 points, with similar cumulative error profiles and preserved predicted-versus-true time structure (Supplementary Fig. 4a). By contrast, the two-time-point setting showed more distorted phase recovery. This limitation was more pronounced downstream: amplitude and phase recovery deteriorated sharply with only two time points, whereas performance improved substantially at four time points and remained comparatively stable with denser sampling (Supplementary Fig. 4b). Rhythm-state assignment showed the same pattern, with substantial module mixing in the two-time-point setting and clearer diagonal recovery from four time points onward (Supplementary Fig. 4c,d). Thus, in this simulation setting, four zeitgeber time points represent a practical lower bound for stable phase-resolved gene-level analysis, whereas two time points are insufficient for reliable parameter estimation and rhythm-state assignment.

We also assessed sensitivity to cell number. Cell-phase inference was relatively stable across the tested range from 20,000 to 600 cells, with similar cumulative error profiles (Supplementary Fig. 5a). In contrast, gene-level rhythmic parameter estimation and rhythm-state assignment were more cell-number dependent. Amplitude and phase recovery were poorest at 600 and 1,500 cells, improved from 3,000 cells and became more stable at higher cell numbers (Supplementary Fig. 5b). Module confusion was most evident at 600 cells and decreased substantially as cell number increased (Supplementary Fig. 5c,d). Runtime analysis showed that phase inference was computationally lightweight, whereas gene-level rhythmic parameter estimation accounted for most of the runtime and increased strongly with cell number (Supplementary Fig. 6). These perturbation benchmarks define practical operating ranges for applying scRhythmBayes to sparse single-cell circadian datasets.

### Periportal and pericentral hepatocytes occupy distinct circadian states, revealing temporal zonation of the liver lobule

We next applied scRhythmBayes to a mouse liver single-cell multiomic dataset to test whether spatially defined hepatocyte populations also differ in circadian organization^10^. After annotation, we focused on peri-central vein hepatocytes (Hep.CV) and peri-portal vein hepatocytes (Hep.PV), two hepatocyte subtypes with established spatial and metabolic specialization (Fig. 4a and Supplementary Fig. 7a)^11,12^. We sought to determine whether this spatial distinction reflects subtype-specific differences in circadian organization.

**Fig. 4.**
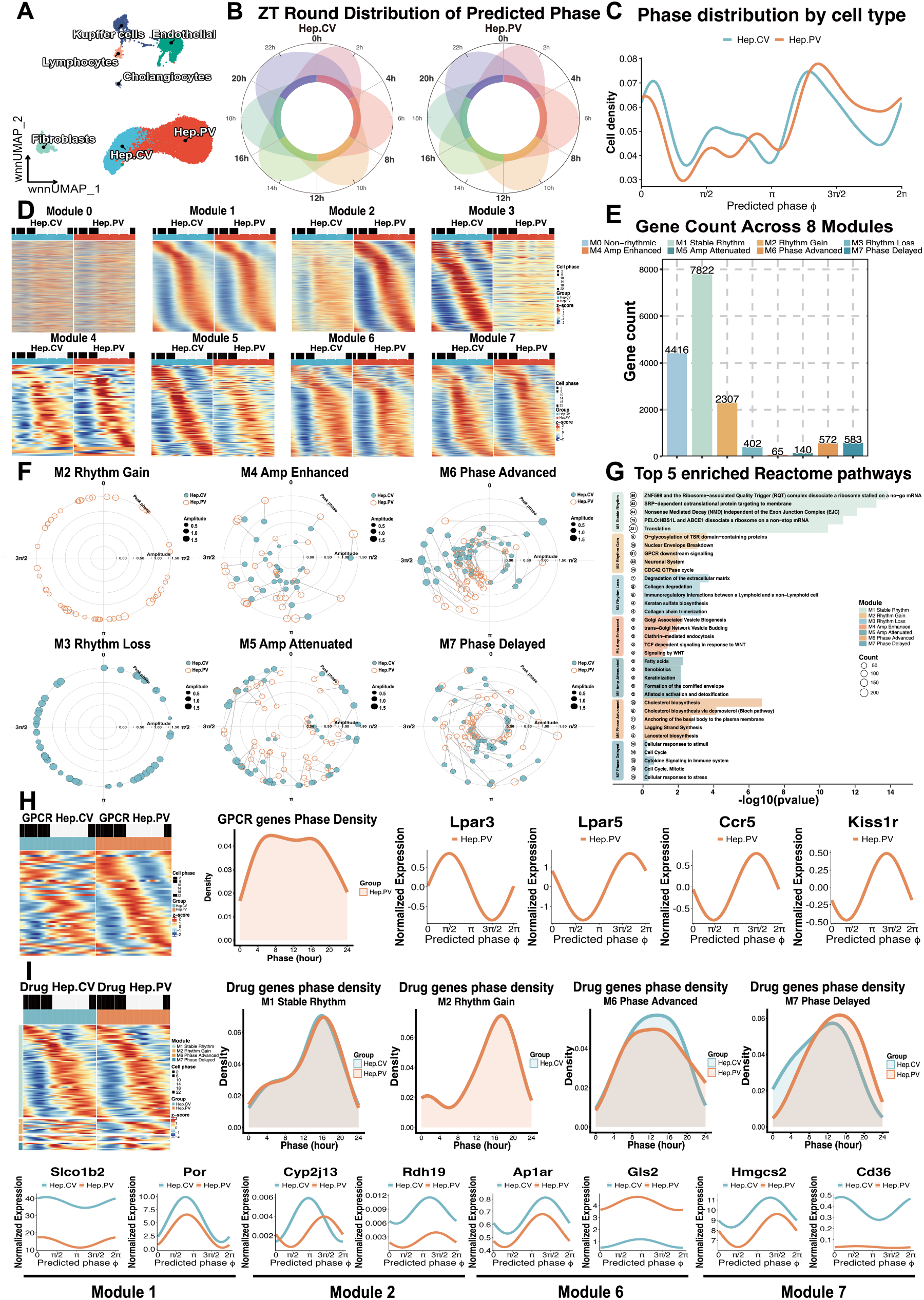
Application of scRhythmBayes to circadian programs in two hepatocyte subtypes. **a**, UMAP of the annotated liver single-cell dataset, showing the major cell populations and the two hepatocyte subtypes analyzed downstream, peri-central vein hepatocytes (Hep.CV) and peri-portal vein hepatocytes (Hep.PV). **b**, Circular summaries of predicted phase distribution across rounded ZT groups in the two hepatocyte subtypes. **c**, Density distributions of predicted phase in the two hepatocyte subtypes. **d**, Paired heatmaps of gene expression patterns across the eight scRhythmBayes modules in the two hepatocyte subtypes. **e**, Bar plot showing gene counts across the eight predefined rhythm-state modules. **f**, Polar plots of representative genes from differential rhythm modules. Angular position represents phase and radial distance represents amplitude, illustrating rhythm gain, rhythm loss, amplitude-enhanced, amplitude-attenuated, phase-advanced, and phase-delayed patterns across the two hepatocyte subtypes. **g**, Top enriched Reactome pathways across rhythm-state modules. **h**, Heatmaps and phase-density summaries of genes from the GPCR downstream signaling pathway, highlighted from Module2 in **g**, showing subtype-specific phase patterns in the two hepatocyte subtypes. **i**, Heatmaps, phase-density summaries, and representative expression trajectories of genes related to drug metabolism, highlighting subtype-specific rhythmic programs across selected modules in the two hepatocyte subtypes.

scRhythmBayes-inferred phase distributions showed that Hep.CV and Hep.PV both covered the 24-h cycle but differed in phase occupancy. Circular summaries across rounded ZT groups and continuous phase-density curves revealed subtype-specific enrichment across different phase intervals (Fig. 4b,c). Additional circular summaries and core clock gene phase plots supported subtype-associated temporal structure in the two hepatocyte populations (Supplementary Fig. 7b–d). These results indicate that Hep.CV and Hep.PV are not only spatially distinct, but also occupy different circadian states within the liver lobule.

We then examined gene-level circadian remodeling between the two hepatocyte subtypes. Paired module heatmaps showed a large stable-rhythm background together with rhythm gain, rhythm loss, amplitude remodeling and phase-shift states (Fig. 4d). Module counts showed that stable rhythm was the largest category (7,822 genes), followed by non-rhythmic genes (4,416 genes); rhythm gain was more frequent than rhythm loss (2,307 versus 402 genes), and phase-shift modules were more abundant than amplitude-remodeling modules (Fig. 4e). Polar plots of representative genes confirmed the expected geometry of the module definitions: gain and loss reflected appearance or disappearance of rhythmic amplitude, amplitude modules showed radius changes at similar phase, and phase modules showed angular displacement at comparable amplitude (Fig. 4f). Module-wise amplitude and phase distributions further supported separation of rhythm-state features between Hep.CV and Hep.PV (Supplementary Fig. 7e).

Reactome enrichment analysis showed that differential rhythm-state modules were associated with functionally coherent programs, including translation, GPCR downstream signaling, extracellular matrix organization, WNT signaling, fatty-acid and xenobiotic metabolism, cholesterol biosynthesis and cellular stress responses (Fig. 4g). We focused on GPCR downstream signaling because this pathway was enriched among rhythm-gain genes and links hepatocytes to extracellular, hormonal and metabolic cues^13^. GPCR-related genes showed subtype-specific phase-density patterns and representative trajectories, indicating that Hep.CV and Hep.PV differ not only in expression level but also in the inferred timing of pathway activity (Fig. 4h).

Drug-metabolism-related genes provided a second example of subtype-specific rhythmic organization, consistent with extensive circadian regulation of hepatic xenobiotic and drug-metabolism pathways^1,14,15^. Heatmaps, phase-density summaries and representative trajectories showed distinct temporal patterns between Hep.CV and Hep.PV across selected rhythm-state modules (Fig. 4i). Genes associated with hepatic uptake, cytochrome P450 activity, oxidation-reduction metabolism, amino-acid metabolism and lipid handling showed subtype-specific rhythmic trajectories. Additional module-associated trajectories supported the same interpretation for drug-metabolism-related programs in Modules 1, 2, 6 and 7 (Supplementary Fig. 8). Together, these results support a temporal dimension of liver zonation. Periportal and pericentral hepatocytes differ in circadian phase occupancy, rhythm-state composition, and pathway timing.

We further evaluated this conclusion in an independent hepatocyte single-cell dataset^12^. Hep.CV and Hep.PV again showed distinct predicted phase-density and circular phase patterns (Supplementary Fig. 9a–c). Module-wise amplitude and phase distributions, core clock gene phase summaries, paired heatmaps, pathway enrichments and representative polar plots were consistent with subtype-associated rhythmic remodeling (Supplementary Fig. 9d,e,g–i). Cross-dataset comparison of rhythm-state module proportions showed concordance between the primary and independent hepatocyte datasets (Supplementary Fig. 9f). Thus, the independent dataset supported the reproducibility of temporal zonation between Hep.CV and Hep.PV, although exact gene-level phase estimates remained sensitive to sampling and dataset composition.

### Cell-type-specific rhythmic programs reveal a temporal dimension of cellular identity

In complex tissues, the most informative pairwise cell-type comparison may not be known in advance. We therefore developed SRB_OVR, a one-versus-rest rhythmic inference procedure, to identify cell-type-specific temporal programs across liver cell types. SRB_OVR uses cell-level circadian phases inferred across all cells as a shared temporal coordinate system. For each target cell type, gene-level rhythmic parameters are then estimated in the target population and in all remaining cell types pooled as Rest. A gene is defined as a cell-type-specific rhythmic marker when it shows rhythmicity with sufficient amplitude in the target cell type but lacks comparable rhythmic evidence in Rest. Thus, SRB_OVR defines cell-type specificity in the temporal domain by testing whether a gene is rhythmic in one target population but not in the pooled remainder (Fig. 5a)^16^.

**Fig. 5.**
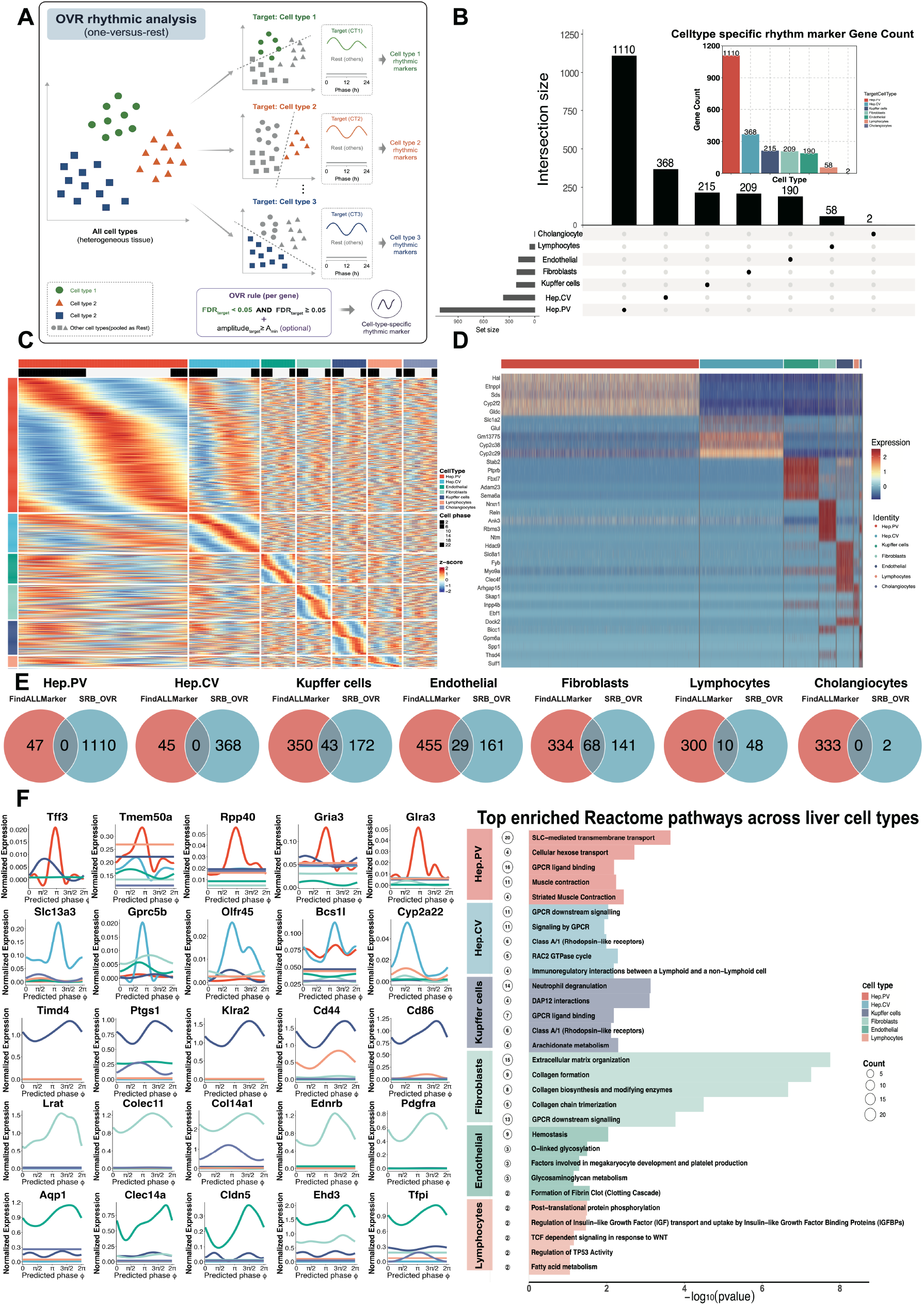
Application of SRB_OVR reveals cell-type-specific rhythmic programs across liver cell types. **a**, Schematic overview of SRB_OVR analysis. After global cell-level phase inference, each target cell type is compared against the pooled Rest background to identify target-specific rhythmic markers whose rhythmic structure is specific to the target cell type rather than to the tissue as a whole. **b**, UpSet plot and marker-count summary showing the overlap structure of SRB_OVR-defined cell-type-specific rhythmic markers across liver cell types. **c**, Phase-ordered heatmap of SRB_OVR-defined rhythmic markers across liver cell types, showing cell-type-resolved dynamic expression programs over predicted circadian phase. **d**, Heatmap of conventional static marker-gene expression across liver cell types identified by Seurat FindAllMarkers. **e**, Venn diagrams comparing SRB_OVR-defined rhythmic markers with FindAllMarkers-derived static markers for each liver cell type. **f**, Representative phase-ordered expression trajectories of SRB_OVR-defined rhythmic marker genes and Reactome pathway enrichment of cell-type-specific rhythmic programs across liver cell types.

SRB_OVR identified cell-type-specific rhythmic marker sets with strong cell-type dependence. Hep.PV contained the largest number of SRB_OVR-defined markers, followed by Hep.CV, Kupffer cells, fibroblasts, endothelial cells, lymphocytes and cholangiocytes (Fig. 5b). Importantly, the overlap structure indicated that most rhythmic markers were restricted to individual liver cell types rather than shared broadly across the tissue. Cell-type-resolved phase distributions and core clock gene summaries supported the presence of distinct temporal organization across major liver populations (Supplementary Fig. 10a–c).

Phase-ordered heatmaps showed that SRB_OVR markers formed dynamic expression programs across predicted circadian phase rather than static cell-type expression blocks (Fig. 5c). We therefore compared these rhythmic markers with conventional static markers identified by Seurat FindAllMarkers, a standard expression-based strategy for single-cell cell-identity annotation^16^. Here, static markers refer to cell-type markers defined by average differential expression across cells, without modeling circadian phase, rhythmic amplitude or phase-dependent expression dynamics. Static marker heatmaps recovered expected cell-identity patterns (Fig. 5d), whereas SRB_OVR captured a distinct temporal layer of cell identity. The limited overlap between rhythmic and static marker sets showed that cell-type-specific rhythmic programs were not simply a consequence of average expression specificity (Fig. 5e). These results indicate that cellular identity is encoded not only by which genes are expressed, but also by when cell-type-specific programs are activated across the circadian cycle. This distinction is consistent with prior work showing that liver transcriptional programs can be jointly organized by lobular position and circadian time^10-12^. Cumulative phase and amplitude summaries further characterized the temporal coverage and rhythmic strength of SRB_OVR-defined markers across liver cell types (Supplementary Fig. 10d,e).

Representative trajectories and Reactome enrichment analyses supported the biological coherence of these temporal marker programs (Fig. 5f). Hepatocyte-associated rhythmic markers were enriched for transport, metabolic and GPCR-related signaling pathways; Kupffer-cell markers were enriched for immune-interaction and inflammatory pathways; fibroblast markers were enriched for extracellular-matrix and collagen-associated programs; and endothelial markers were enriched for hemostasis and glycosylation-related pathways. These results show that SRB_OVR resolves cell-type-specific temporal programs and reveals a temporal dimension of cellular identity that is largely invisible to conventional static marker analysis.

### Cell-phase coordination and DTW reveal cell-type-selective temporal disorganization in human heart disease

Finally, we applied scRhythmBayes to a human heart disease dataset to determine whether circadian disruption occurs uniformly across the tissue or selectively affects specific cardiac cell populations^17^. We focused on cell-phase organization as an initial measure of temporal vulnerability across cardiac cell types. We therefore summarized inferred phases using two complementary measures. The mean resultant length, ρ, was computed within each biological sample and cell type to quantify phase coordination. Lower ρ indicates weaker concentration of cells around a common inferred phase. Dynamic time warping (DTW) was used to compare normalized phase-distribution profiles between normal circadian rhythm (NCR) and abnormal circadian rhythm (ACR). This analysis captures shifts, broadening, and misalignment of phase occupancy (Fig. 6a).

**Fig. 6.**
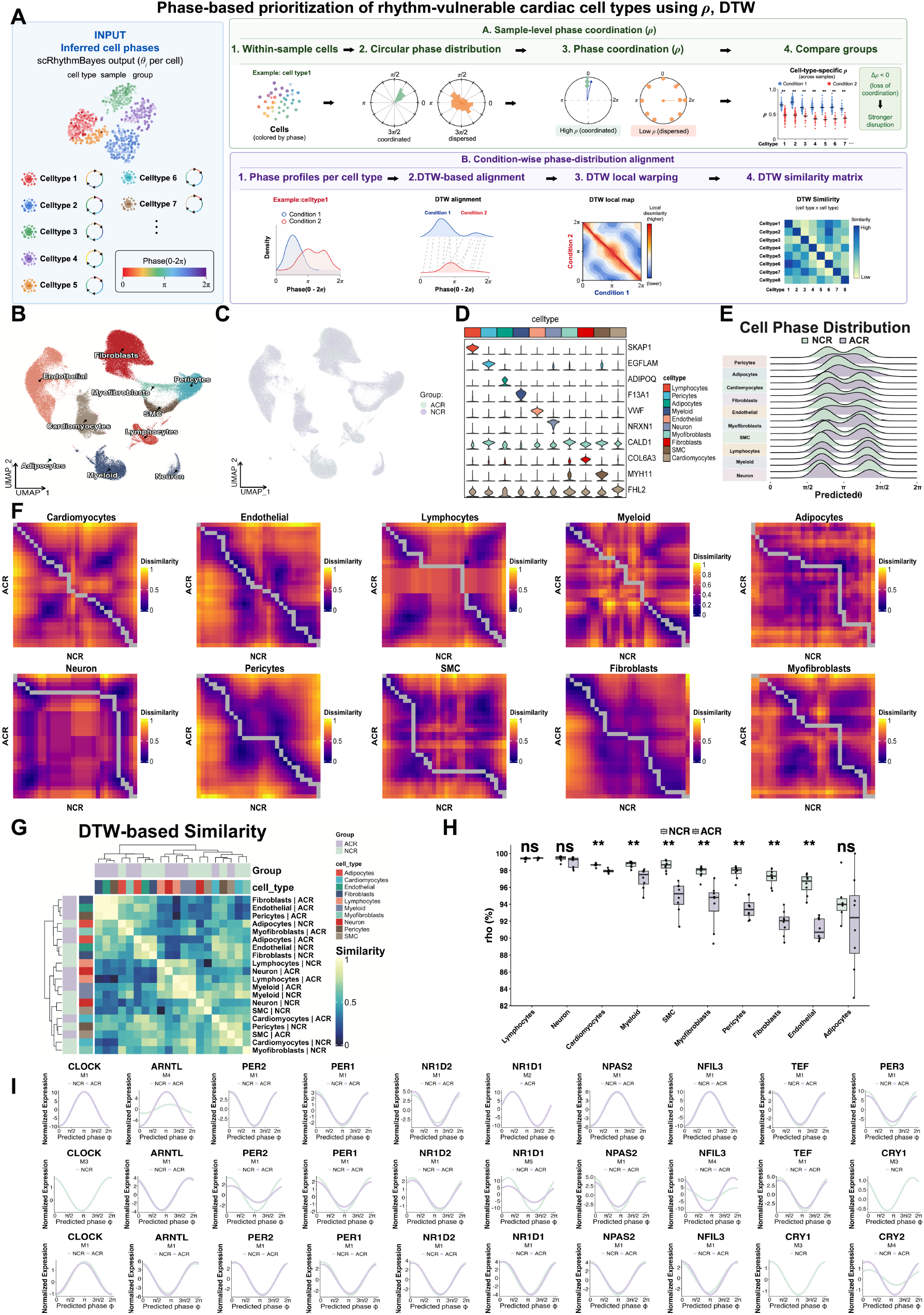
Application of cell-phase coordination and DTW-based alignment to prioritize rhythm-vulnerable cell types in the diseased human heart. **a**, Computational workflow for cell-phase-level prioritization using sample-level phase coordination and DTW-based phase-distribution alignment. Inferred cell phases are organized by cell type, sample and group. Within-sample phase coordination is quantified by the mean resultant length, ρ, and compared between conditions across biological samples. Condition-specific phase distributions are then converted into normalized phase profiles and compared using dynamic time warping to quantify between-condition phase-profile divergence. **b**, UMAP of the annotated human heart single-cell dataset, showing the major cardiac cell populations analyzed downstream. **c**, UMAP overlaid with scRhythmBayes-inferred cell phases. **d**, Marker-gene violin plots supporting the major cardiac cell-type annotations. **e**, Cell-phase distributions across major cardiac cell types in normal circadian rhythm and abnormal circadian rhythm groups. **f**, DTW-based phase-profile dissimilarity maps comparing normalized NCR and ACR phase distributions for each cardiac cell type. **g**, DTW-based similarity heatmap summarizing phase-distribution alignment across cardiac cell-type and condition profiles. **h**, Cell-type-wise comparison of sample-level mean resultant length, ρ, between NCR and ACR. Each point represents a biological sample within a cell type. **i**, Phase-ordered expression trajectories of representative core clock and cell-type-relevant genes in prioritized cardiac cell types, including neurons, smooth muscle cells and cardiomyocytes. Additional cardiac cell types are shown in Supplementary Fig. 11.

After annotation of major cardiac cell populations, including cardiomyocytes, fibroblasts, endothelial cells, myofibroblasts, pericytes, smooth muscle cells, myeloid cells, lymphocytes, neurons and adipocytes, inferred phases could be projected back onto the heart atlas (Fig. 6b,c)^17^. Marker-gene distributions supported the major cell-type annotations (Fig. 6d). Cell-phase distributions across normal circadian rhythm (NCR) and abnormal circadian rhythm (ACR) groups showed that some cell types retained similar phase profiles, whereas others displayed altered phase occupancy or broader distributional distortion (Fig. 6e).

DTW-based phase-profile dissimilarity maps and similarity summaries provided a global view of between-condition phase-distribution alignment across cardiac cell types (Fig. 6f,g). In parallel, sample-level ρ was computed within each cell type and biological sample before group comparison. Across multiple cardiac populations, NCR samples showed higher phase coordination than ACR samples. This indicates that abnormal circadian rhythm is associated with cell-type-selective loss of temporal synchrony rather than a uniform disruption across the heart (Fig. 6h). Cell-type-resolved phase distributions and sample-level ρ comparisons for additional cardiac populations are shown in Supplementary Fig. 11a,b.

We selected neurons, smooth muscle cells, and cardiomyocytes for main-text visualization because they represent distinct axes of cardiac temporal regulation. These include neuro-cardiac timing, vascular rhythmicity, and intrinsic myocardial clock function^18-21^. Neurons showed altered DTW-based alignment, consistent with a possible shift in neuro-cardiac timing. Smooth muscle cells showed reduced phase coordination, suggesting altered vascular timing. Cardiomyocytes represent the main contractile and electrophysiological compartment, and the source DCM study reported reduced ventricular mechanical strength in the ACR group^17^. Core clock and cell-type-relevant genes in these populations showed condition-associated phase-pattern differences between NCR and ACR (Fig. 6i). Additional cardiac cell types showed phase-ordered gene trajectories in the supplementary analysis (Supplementary Fig. 11c). These results indicate that human heart disease is associated with cell-type-selective temporal disorganization across myocardial, vascular, and neuro-cardiac compartments^17^.

## Discussion

scRhythmBayes treats single-cell circadian analysis as a distinct statistical problem rather than a direct extension of bulk time-series analysis. Existing methods have provided important solutions for cyclic ordering, cell-phase inference, rhythmicity detection, waveform estimation, and differential rhythmicity^3–9^. However, these tasks are usually formulated under different assumptions. In particular, gene-level models often assume that time is already known, whereas cell-phase methods may stop after ordering or phase prediction. scRhythmBayes introduces dedicated probabilistic models for latent cell phase, phase-confidence-aware gene-level inference, and differential rhythm-state classification. Sample-level zeitgeber time is used only to orient the circular phase space. It is not treated as the true phase of every cell. This distinction is important because cells collected at the same zeitgeber time may differ in circadian state owing to cell type, local environment, biological condition, or incomplete synchronization. The resulting phase coordinate therefore represents a latent dimension of cellular state rather than a relabeling of sample collection time.

The rhythm-state model also illustrates an important trade-off between biological continuity and interpretability. Circadian remodeling does not necessarily occur as sharply separated categories. Stable rhythmicity, amplitude attenuation, and rhythm loss can form a continuum. This helps explain why the amplitude-attenuated state, M5, is more difficult to classify than the stable-rhythm state, M1. M1 is supported when both conditions retain clear rhythmicity with similar amplitude and phase. It therefore occupies a relatively broad and well-separated region of parameter space. By contrast, M5 lies close to several boundaries. A modest reduction in amplitude may resemble stable rhythmicity, whereas a stronger reduction may approach rhythm loss or non-rhythmicity. Sparse counts, low expression, phase uncertainty, and limited cell number can move a gene across these boundaries. The eight modules should therefore be interpreted as dominant summaries of remodeling rather than absolute biological classes. Posterior probabilities and assignment margins are particularly important for genes with mixed amplitude and phase changes. They allow borderline patterns to remain visible instead of being hidden by a single hard label.

The liver analysis suggests that classical lobular zonation may include a temporal component. Periportal and pericentral hepatocytes are exposed to different oxygen levels, nutrients, hormones, and metabolic substrates, and they perform distinct physiological functions^10–12^. These spatial gradients may also alter the timing at which cellular programs are activated. In this context, the predominance of rhythm gain over rhythm loss is more consistent with subtype-specific recruitment of rhythmic programs than with a simple erosion of circadian regulation. The relative abundance of phase-shift states further suggests that temporal repositioning may be at least as important as changes in rhythmic strength. Two hepatocyte populations may therefore execute related functions at different times of day, even when their average expression levels are similar. This interpretation is relevant to GPCR signaling, lipid handling, xenobiotic metabolism, and drug-processing pathways^1,13–15,22–24^. It raises the possibility that spatial zonation and circadian timing jointly shape hepatic pharmacology. The independent dataset supports the broader concept of temporal zonation, although exact gene-level phase estimates remain sensitive to sampling density and dataset composition.

The SRB_OVR and heart analyses extend this temporal view from tissue zonation to cellular identity and disease vulnerability. Conventional marker analysis defines cell identity through average expression differences^16^. A rhythmic marker instead captures whether a gene follows a cell-type-specific temporal program. These two forms of specificity are not equivalent. A gene may lack strong average-expression specificity but still show a distinctive phase or amplitude pattern in one cell population. Cellular identity may therefore be encoded by both expression level and temporal organization. SRB_OVR should nevertheless be interpreted as a discovery procedure. Pooling all non-target cells into Rest can dilute rhythmic signals shared by several minority populations, so target specificity requires independent validation. A related issue arises in the heart. Reduced phase coordination does not necessarily indicate complete clock failure, and altered DTW alignment does not establish a causal mechanism. These patterns may reflect desynchronization, phase redistribution, altered cell-state composition, or differential responses to disease. The value of the analysis lies in identifying candidate cellular compartments for follow-up. Neuronal, vascular, and myocardial populations represent distinct axes through which circadian disruption may influence cardiac physiology^17–21,25,26^. This supports a model in which disease-associated temporal disruption is cell-type selective rather than uniform across the tissue.

Several limitations define the current scope of scRhythmBayes. Direct ground truth for cell-level circadian phase is rarely available in real tissues. Phase recovery must therefore be evaluated through simulations, known clock-gene behavior, cross-dataset consistency, and biological plausibility. The gene-level model uses posterior phase summaries and confidence weights rather than fully marginalizing over the complete phase posterior. This approximation improves computational scalability but means that downstream uncertainty is conditional on the inferred phase summaries. The method also requires informative seed genes, sufficient temporal coverage, and adequate cell numbers within each stratum. Sparse experimental designs may support broad phase-level prioritization but remain insufficient for reliable gene-level remodeling analysis. Future studies should combine scRhythmBayes with denser temporal sampling, independent cohorts, perturbation experiments, and spatially resolved measurements. Integration with chromatin accessibility, proteomic, and metabolomic data may further clarify how temporal programs are established and propagated across molecular layers. More broadly, scRhythmBayes supports a view of single-cell state that includes cell type, biological condition, spatial context, and circadian time.

## Methods

### Overview of scRhythmBayes

scRhythmBayes is a hierarchical probabilistic algorithm for circadian analysis of heterogeneous single-cell transcriptomic data. The input consists of a raw gene-by-cell count matrix, cell-level metadata containing sample-level zeitgeber time, and one or more stratum labels such as cell type, condition, or sample. The algorithm estimates latent cell-level circadian phase, gene-level rhythmic parameters, and interpretable differential rhythm states.

The algorithm contains three statistically coupled inference components. First, cell-level circadian phase is inferred from rhythmic seed genes using a seed-gene count likelihood and a stratum-aware circular mixture prior centered on sample-level zeitgeber time. Second, conditional on the inferred phases and cell-level phase support, gene-level rhythmic parameters are estimated within each biological group or stratum. These parameters include mesor, mean expression, amplitude, phase, overdispersion, dropout rate, and rhythmicity significance. Third, for two-group comparisons, genes are assigned posterior support across predefined rhythm-state categories using group-specific rhythmic evidence, amplitudes, and phases.

Depending on the analysis objective, scRhythmBayes can be used for cell-phase inference alone, gene-level rhythmic modeling, or full rhythm-state remodeling analysis. In heterogeneous tissues, the inferred phases and gene-level parameters also support one-versus-rest rhythmic marker discovery. In multi-condition datasets, inferred cell phases can be summarized using sample-level phase coordination and dynamic time warping-based phase-distribution alignment to prioritize cell types with condition-associated temporal disorganization.

### Cell-level phase inference from seed rhythmic genes

#### Input data and notation

Let 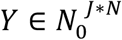 denote the raw count matrix of *J* selected seed genes across *N* cells, where *Y*_*ji*_ is the raw count of seed gene *j* in cell *i* . Each cell *i* is associated with a sample-level zeitgeber time *ZT*_*i*_, a stratum label *c*(*i*), and a library-size offset *L*_*i*_, computed from the raw count matrix. Seed genes are either supplied by the user or selected from a built-in rhythmic seed panel.

For each cell *i*, scRhythmBayes introduces a latent circadian phase

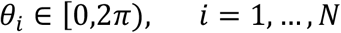

and denotes the full collection of cell phases by

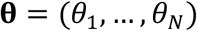

The objective of this layer is to infer the posterior distribution of **θ** from the seed-gene count matrix together with prior temporal information. Sample-level zeitgeber time is mapped to a circular anchor

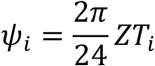

which is treated as prior information about the phase of cell *i*, rather than as its true circadian phase.

#### Seed-gene count likelihood

For seed gene *j* in cell *i*, counts are modeled as

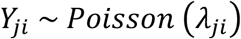

with log-mean

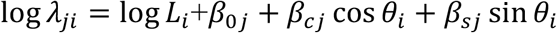

Here, **β**_0*j*_ is a gene-specific intercept, whereas *β*_*c j*_ and *β*_*s j*_ are the cosine and sine coefficients of seed gene *j*, respectively. Under this formulation, all seed genes within a given cell share the same latent phase *θ*_*i*_, while each gene retains its own baseline and rhythmic response. The joint seed-gene likelihood therefore defines, for each cell, a phase-dependent expression-based evidence surface over the circadian circle.

#### Stratum-aware circular phase prior

To accommodate heterogeneous single-cell data, in which cells collected at the same sample-level zeitgeber time need not share the same true phase and different strata may exhibit systematic phase offsets and distinct levels of temporal concentration, scRhythmBayes specifies a stratum-aware circular mixture prior over *θ*_*i*_ :

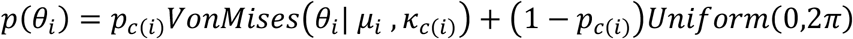

where

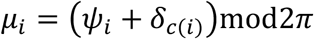

In this model, *δ*_*c*_ denotes a stratum-specific phase shift relative to the sample-level anchor, *κ*_*c*_ is a stratum-specific concentration parameter, and *p*_*c*_ is the mixing weight of the structured rhythmic component. The uniform component allows cells with weak rhythmic evidence to retain diffuse posterior support rather than being forced toward the zeitgeber anchor.

In the default implementation, the initial von Mises concentration is *κ*_*c*_ = 5 the initial mixture weight is *p*_*c*_ = 0.85, and the minimum number of cells required for stratum-level prior updating is 200. During iterative updating, *κ*_*c*_ is constrained to the interval [0.3,50], and *p*_*c*_ is constrained to [0.05,0.995] for numerical stability.

#### Discrete posterior approximation

Because the posterior distribution of *θ*_*i*_ under the seed-gene likelihood and stratum-aware circular prior is not analytically tractable, scRhythmBayes approximates it on a discrete phase grid {ϑ_1_, …, ϑ_*M*_} spanning [0,2*π*), where ϑ_*m*_ denotes the phase value at grid point *m*.

For each seed gene *j* and phase grid point ϑ_*m*_, the phase-specific log-intensity is defined as

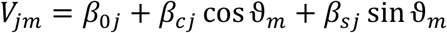

Given the current seed-gene coefficients, let *l*_*im*_ denote the seed-gene joint log-likelihood of cell *i* when its latent phase is fixed at the grid point ϑ_*m*_. Then

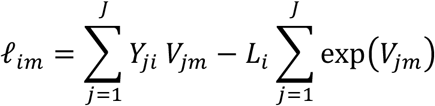

up to an additive constant independent of ϑ_*m*_. Thus, *l*_*im*_ is the likelihood score of cell *i* evaluated at the candidate phase ϑ_*m*_, and defines a phase-dependent likelihood surface over the circular grid for each cell.

For cell *i*, the discrete prior mass at grid point ϑ_*m*_ is defined as

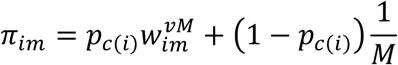

where

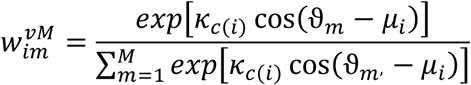

The posterior phase distribution of cell *i* is then approximated on the grid by

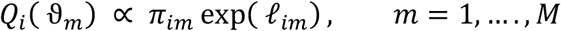

Posterior expectations of the circular basis functions are computed as

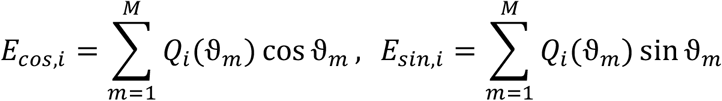

and are used to refit the seed-gene Poisson coefficients (*β*_0*j*_, *β*_*cj*_, *β*_*sj*_ ). The updated coefficients are then used to recompute the cell-specific phase posterior.

To quantify whether a cell *i* is better supported by the structured von Mises component than by the circular uniform component, scRhythmBayes computes a cell-level structured-component responsibility

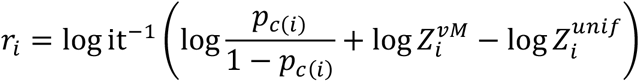

where

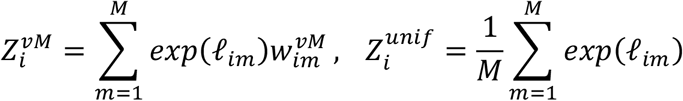

Thus, *r*_*i*_ is interpreted as a posterior responsibility for the structured component, rather than as an ad hoc score. Cells with weaker support are adaptively shrunk toward a flatter posterior, and stratum-specific prior parameters ( *δ*_*c*_, *κ*_*c*_, *p*_*c*_ ) are updated from responsibility-weighted circular summaries. These updates are iterated for a fixed number of rounds.

The final cell phase is summarized by the posterior mean direction

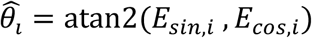

which is converted to circadian time in hours as

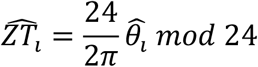

scRhythmBayes additionally reports the posterior resultant length

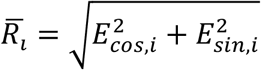

as a measure of phase concentration, together with the cell-level responsibility *r*_*i*_. The output of this layer is therefore the posterior phase summary (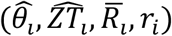, *r*_*i*_), which is passed to downstream gene-level rhythmic modeling.

### Gene-level rhythmic parameter estimation conditional on inferred phase

#### Input to gene-level modeling

Given the inferred cell phases 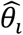 and cell-level structured-component responsibilities *r*_*i*_ from the cell-level inference layer, scRhythmBayes next fits gene-level rhythmic models to the raw count matrix

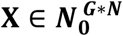

where *X*_*gi*_ denotes the raw count of gene *g* in cell *i*, with *g* = 1, …, *G*. Each cell is associated with a biological condition, and gene-level rhythmic parameters are estimated separately within each condition.

To account for library-size variation, each cell *i* is assigned a size factor *s*_*i*_, estimated from the sparse count matrix using a median-ratio normalization procedure. The corresponding log-size factor log *s*_*i*_ is used as an offset in all downstream gene-level models.

The cell-level responsibility *r*_*i*_ is propagated to the gene-level layer through a bounded numerical weight *p*_*i*_, which is used in downstream mixture-likelihood optimization. In the default implementation,

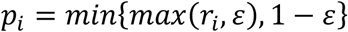

where *ε* is a small numerical constant used only for optimization stability. Thus, the quantity passed between layers is the phase-layer structured-component responsibility, with mild numerical bounding applied in implementation.

#### Weighted gene-level rhythmic likelihood

For a given gene *g* within a given condition, scRhythmBayes compares a rhythmic model against a nested non-rhythmic null model. Under the full rhythmic model, counts are assumed to follow a zero-inflated negative binomial distribution,

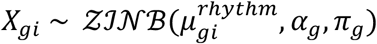

with mean

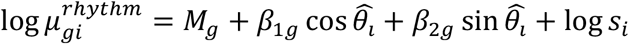

Here, *M*_*g*_ is the log-scale baseline expression, *β*_1*g*_ and *β*_2*g*_ are the cosine and sine coefficients, *α*_*g*_ is the negative-binomial overdispersion parameter, and *π*_*g*_ is the zero-inflation probability.

The corresponding null model removes the phase-dependent terms,

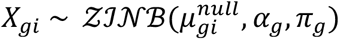

with

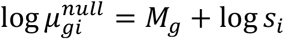

Gene-level rhythmic modeling uses the posterior mean phase 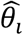 and the structured-component responsibility *r*_*i*_ from the cell-level phase-inference layer. The posterior mean phase provides the inferred circadian position of each cell, whereas *r*_*i*_, represented in optimization by the bounded weight *p*_*i*_, controls how strongly each cell contributes to rhythmic parameter estimation. This design incorporates cell-level phase confidence into gene-level fitting while maintaining scalability for genome-wide single-cell analysis.

Let

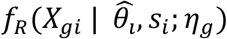

denote the single-cell probability mass under the rhythmic model, and let

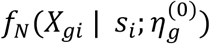

denote the corresponding single-cell probability mass under the null model, where *η*_*g*_ and 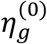 collect the gene-level parameters under the full and null models, respectively. The gene-level log-likelihood is then defined as

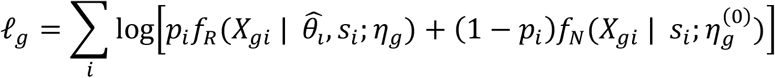

Thus, cells with stronger phase support contribute more strongly to the rhythmic component of the likelihood, whereas weakly supported cells contribute relatively more to the null component.

#### Parameter estimation and rhythmicity testing

For each gene within each condition, parameters (*M*_*g*_, *β*_1*g*_, *β*_2*g*_, log *α*_*g*_, logit(*π*_*g*_)) are estimated by likelihood optimization. Initial values for *M*_*g*_, *β*_1*g*_, and *β*_2*g*_ are obtained from a weighted linear regression of log-transformed normalized expression on 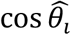 and 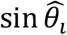, using the cell-level weights *p*_*i*_. Initial values for *α*_*g*_ and *π*_*g*_ are derived from the empirical mean-variance relationship and zero fraction, respectively.

From the fitted cosine-sine coefficients, gene-level rhythmic summaries are defined as

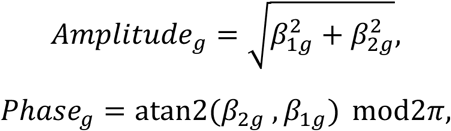

and

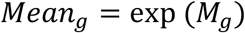

The fitted *M*_*g*_ is also retained as the log-scale baseline (or log-mesor) parameter, together with the overdispersion parameter *α*_*g*_ and the dropout rate *π*_*g*_.

Rhythmicity is assessed by likelihood-ratio testing between the full and null models,

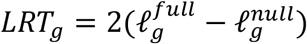

where 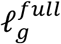 and 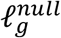 are the optimized log-likelihoods under the full and null models, respectively. Because the full model introduces two additional phase-dependent parameters (*β*_1*g*_, *β*_2*g*_), P-values are computed using a *χ*^2^ reference distribution with 2 degrees of freedom and adjusted across genes within condition using the Benjamini–Hochberg procedure.

#### Group-wise parameter summaries

After fitting each gene separately within each condition, scRhythmBayes stores the resulting estimates in both long and wide formats. For each gene-condition pair, the model reports log-scale baseline, mean, amplitude, phase, overdispersion, dropout rate, likelihood-ratio statistic, nominal P-value, and false discovery rate.

For two-condition analyses, cross-condition contrasts are additionally summarized as

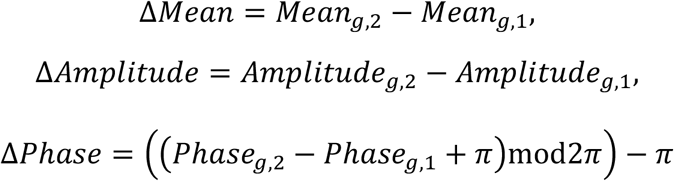

These quantities summarize cross-condition rhythmic differences. For downstream differential rhythm state classification, amplitude and phase contrasts are further transformed into module-classification features in the next layer. When analyses are performed separately within a given cell type or user-defined stratum, these group-wise summaries additionally support identification of cell-type-specific rhythmic gain, rhythmic loss, amplitude modulation, and phase shift across conditions.

#### Bootstrap uncertainty quantification

To quantify estimation uncertainty for the most informative genes, scRhythmBayes performs within-condition bootstrap resampling on a selected subset of genes prioritized by rhythmic significance and amplitude. For each selected gene, cells are resampled with replacement, the gene-level model is refit, and bootstrap estimates of mean, amplitude, and phase are collected.

Confidence intervals for mean and amplitude are computed from empirical quantiles of the bootstrap distribution. Because phase is circular, bootstrap phase intervals are computed on the wrapped difference scale,

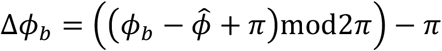

where 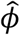 is the point estimate and *ϕ*_*b*_ is the bootstrap estimate. Quantile limits on Δ*ϕ*_*b*_ are then mapped back to the circular scale to obtain the final phase interval. This preserves the circular geometry of phase uncertainty.

### Differential rhythm state classification

#### Input to module classification

The module-classification layer takes as input the wide-format gene-level parameter table produced by the previous layer. This layer is formulated for pairwise group contrasts.For each gene g, the required inputs are the group-specific false discovery rates, amplitudes, and phases,

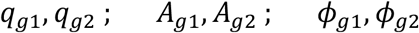

From these quantities, scRhythmBayes derives a set of differential rhythmic features for downstream probabilistic assignment of rhythm states.

#### Feature construction

To quantify cross-group rhythmic differences, the circular phase difference is defined as

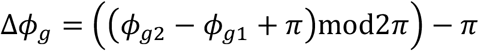

which wraps phase differences to the interval (-*π, π*]. The log-amplitude ratio is defined as

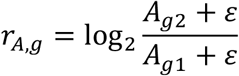

where *ε* is a small constant to avoid division by zero. To encode phase difference on a continuous Euclidean scale, scRhythmBayes further uses the circular embedding

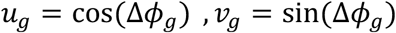

The resulting module feature vector for gene *g* is constructed from

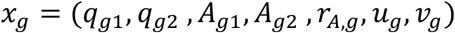

Before probabilistic scoring, these features are centered and scaled across genes.

#### Seed-rule initialization of candidate states

Because the eight differential-rhythm modules represent predefined biological states rather than unconstrained latent clusters, threshold-based seeding is used to anchor module semantics before probabilistic refinement. Final module labels are therefore determined by seed-informed probabilistic scoring in continuous feature space, rather than by threshold rules alone.

scRhythmBayes first assigns each gene a provisional seed state using threshold-based rules on rhythmic significance, amplitude, amplitude change, and phase shift. The thresholds are:

*τ*_*q*_: significance cutoff for rhythmicity,

*τ*_*A*_: minimum amplitude for meaningful rhythmic signal,

*τ*_*dA*_: minimum absolute log-amplitude ratio for amplitude-change modules,

*τ*_*dϕ*_: minimum absolute phase difference for phase-shift modules.

Using these thresholds, genes are assigned to one of eight candidate seed modules: module 0: non-rhythmic,

module 1: stable rhythm,

module 2: rhythm gain,

module 3: rhythm loss,

module 4: amplitude-enhanced rhythmic,

module 5: amplitude-attenuated rhythmic,

module 6: phase-advanced rhythmic,

module 7: phase-delayed rhythmic.

In particular, module 0 includes both genes that are non-significant in both groups and genes that are significant in both groups but have very small amplitudes in both groups. Modules 2 and 3 are defined by asymmetric rhythmic significance across the two groups. Modules 4–7 are defined among genes rhythmic in both groups, using the magnitude of amplitude change and phase shift. When both amplitude and phase differences exceed their respective thresholds, scRhythmBayes resolves the conflict by comparing the standardized scores

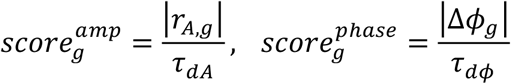

and assigns the gene to the amplitude-dominant or phase-dominant module accordingly.

#### Seed-informed probabilistic module scoring

The provisional seed assignments are then used to construct module-specific probabilistic scoring models in feature space. Let *x*_*g*_ denote the standardized feature vector of gene *g*. For each module *k ∈* {0, …,7}, scRhythmBayes models

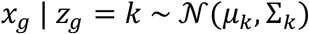

where *z*_*g*_ is the latent module label of gene *g*. The module-specific mean vector *µ*_*k*_ and covariance matrix Σ_*k*_ are estimated from the seed genes currently assigned to module *k*, with shrinkage toward the global feature mean and covariance for numerical stability. Additional diagonal shrinkage is applied to the covariance matrix to reduce ill-conditioning.

A seed-informed module prior is defined as

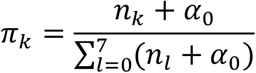

where *n*_*k*_ is the number of seed genes assigned to module *k*, and *α*_0_ is a smoothing constant. For each gene, the module score under module *k* is computed as

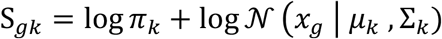

These scores are normalized across modules using a row-wise softmax,

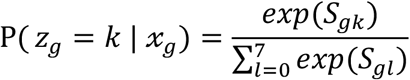

#### Final module assignment and confidence summaries

The final module label of gene *g* is assigned by maximum a posteriori probability,

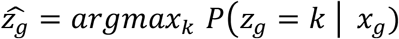

For each gene, scRhythmBayes additionally records the maximum posterior probability

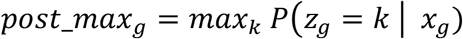

and the margin between the top two posterior probabilities,

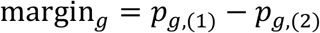

where *p*_*g*,(1)_ and *p*_*g*,(2)_ denote the largest and second-largest module posterior probabilities, respectively. These quantities summarize assignment confidence and separation between competing module labels.

The final output of this layer consists of a gene-level module table containing the group-specific rhythmic features, provisional seed module, final module assignment, posterior module probabilities, and confidence summaries, together with a module-level frequency summary across all genes. The current implementation of the module-classification layer is defined for two-group contrasts. In multi-cell-type settings, target-specific rhythmic programs can additionally be assessed through one-versus-rest comparisons based on cell-type-resolved gene-level summaries.

To assess the dependence of module assignment on seed thresholds, module classification was repeated across a grid of *τ*_*q*_, *τ*_*A*_, *τ*_*dA*_, and *τ*_*dϕ*_ values, and downstream analyses quantified assignment consistency, posterior stability, and preservation of module-level biological conclusions.

#### One-versus-rest extension for multi-cell-type analysis

In heterogeneous tissues containing multiple cell types, scRhythmBayes can be extended through a one-versus-rest (OVR) strategy applied after global phase inference. In this setting, cell-level circadian phase is first inferred jointly across all cells using the seed-gene likelihood together with a stratum-aware circular prior, where the stratum label may correspond to cell type or another user-defined partition. This yields a shared phase summary for all cells while retaining stratum-specific temporal structure.

For downstream gene-level analysis, rhythmic parameters may then be estimated separately within each cell type, yielding cell-type-specific rhythmic summaries. In addition, for a target cell type *c*^*^, an OVR contrast can be constructed by assigning cells from*c*^*^ to the target group and pooling all remaining cell types into a Rest group. The standard two-group gene-level rhythmic modeling procedure is then applied to this target-versus-rest comparison using the same inferred phase summaries from the upstream phase-inference layer. Repeating this procedure across target cell types yields a collection of cell-type-resolved OVR rhythmic summaries.

This OVR strategy provides a practical extension for heterogeneous datasets and enables identification of target-specific rhythmic programs. A gene is labeled as a cell-type-specific rhythmic marker when it is rhythmic in the target cell type but not in the pooled Rest background, optionally together with a minimum-amplitude requirement in the target cell type. Conceptually, this strategy is analogous to one-versus-rest marker analysis such as FindAllMarkers, except that the target of inference is rhythmic structure rather than static expression difference. Thus, scRhythmBayes supports not only repeated two-group analyses within individual cell types, but also OVR-based detection of target-specific rhythmic programs across multiple cell types. In the main text, these genes are referred to as SRB_OVR-defined cell-type-specific rhythmic markers when individual genes are emphasized, and as cell-type-specific rhythmic programs when pathway-level or functional interpretation is emphasized.

### Sample-level phase coordination and DTW-based phase-distribution alignment for multi-condition prioritization

In multi-cell-type datasets containing multiple biological conditions, scRhythmBayes can additionally be used to prioritize cell types for downstream rhythmic comparison based on inferred phase organization. This extension is particularly useful when the primary question is not immediately which genes differ between conditions, but which cell types show the strongest condition-associated perturbation in circadian structure.

For a given cell type *c*, let 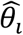 denote the inferred phase of cell *i*, and let each cell additionally be associated with a biological sample label *s*_*i*_ and a group label *g*_*i*_. To quantify within-sample phase coordination, scRhythmBayes computes the mean resultant length separately for each cell type, sample and group combination. For sample s within cell type c, this quantity is defined as

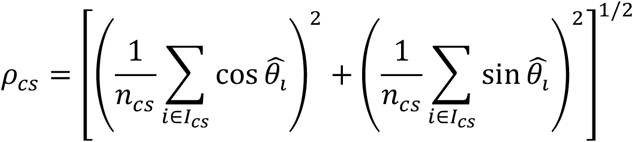

where *I*_*cs*_ is the set of cells belonging to cell type c and sample s, and *n*_*cs*_ is the corresponding number of cells. Thus, *ρ*_*cs*_ summarizes the degree of phase concentration within each biological replicate rather than after pooling all cells across samples. Group-wise comparisons of phase coordination are then performed across samples within each cell type, for example using a rank-based test, to assess whether one condition exhibits systematically weaker within-sample temporal organization than another.

To compare phase-distribution structure between conditions beyond a single concentration summary, inferred phases are converted into cell type– and group-specific phase histograms or density profiles on the circular 24-h domain. These profiles are normalized before alignment. Dynamic time warping (DTW) is then used to quantify the alignment cost or similarity between condition-specific phase profiles. In this setting, DTW provides a cell-type-level summary of between-condition phase-distribution divergence, capturing shifts, broadening or warping of the phase profile that may not be reflected by ρ alone.

Together, sample-level ρ and DTW-based phase-distribution alignment provide complementary summaries of temporal organization. Sample-level ρ quantifies within-sample phase coordination, whereas DTW quantifies between-condition phase-profile alignment. Cell types showing reduced phase coordination, strong between-condition phase-distribution divergence, or both can be prioritized for downstream gene-level rhythmic modeling, representative trajectory visualization or perturbational follow-up.

### Simulation design

We note that the simulation framework is partially matched to the modeling assumptions of scRhythmBayes, because fully realistic simulators with known circadian ground truth are currently unavailable for this problem. Accordingly, these simulations should be interpreted as controlled calibration experiments rather than fully assumption-free benchmarks. Generalizability was therefore further assessed through real-data analyses and downstream evaluations.

### Simulation framework for cell-level phase inference

To evaluate the cell-level phase-inference component, we generated heterogeneous single-cell circadian count data with known sample-level zeitgeber time, cell-level true phase, and cell-type-specific rhythmic structure. The simulated gene universe was initialized from a 4,000-gene set derived from GSE11923, and the rhythmic seed panel was taken from the curated seed-gene set used by the scRhythmBayes phase-inference framework.

Cells were simulated at six symbolic Zeitgeber times (ZT=0,4,8,12,16,20). At each symbolic time point, individual cells were assigned continuous ground-truth times sampled from a normal distribution centered at the corresponding sample-level zeitgeber time with standard deviation *σ*_*t*_ = 0.8*h*, followed by circular wrapping to the 24-h cycle. This design captures within-time-point temporal dispersion while preserving the experimentally observed sample-time anchor. Each time point contained 20,000 cells, equally divided across four cell types.

Cell-specific library sizes were sampled from a log_10_ *normal* distribution with mean log_10_(5000) and standard deviation 0.5. At the gene level, baseline expression, rhythmic amplitude, and phase were specified separately for each gene and cell type. A gene-level baseline was first sampled from a log-normal distribution and then scaled by cell-type-specific drift factors. To mimic marker-gene structure, subsets of non-seed genes were additionally assigned elevated baseline expression in specific cell types. Gene phases were sampled uniformly on the 24-h cycle and then shifted by cell-type-specific phase offsets. Seed genes were forced to remain rhythmic across cell types, whereas non-seed genes were assigned rhythmicity probabilistically, with amplitudes sampled from gamma distributions and truncated to avoid extreme values.

Given the true cell time and gene-by-cell-type parameters, the expected expression template was defined as

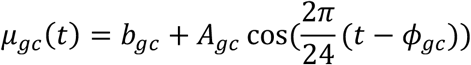

where *b*_*gc*_, *A*_*gc*_, and *ϕ*_*gc*_denote the baseline, amplitude, and phase of gene *g* in cell type *c*, respectively. For each cell, these values were converted to gene-expression proportions, scaled by the cell-specific library size, and sampled using a Poisson model. Expression-dependent dropout was then applied using a logistic function of the expected count intensity. The resulting datasets contain known truth for sample-level zeitgeber time, cell-level true phase, and cell-type-specific rhythmic structure, and were used to benchmark the seed-guided phase-inference layer of scRhythmBayes.

For the phase-inference layer specifically, the seed panel was matched to the seed-guided modeling framework. Robustness to seed mismatch, prior misspecification, and temporal-label perturbation was evaluated separately in downstream stress-test analyses.

### Simulation framework for gene-level rhythmic modeling and differential rhythm state classification

To evaluate the gene-level rhythmic modeling and differential rhythm state-classification layers, we generated synthetic two-group single-cell count data with known group-specific rhythmic parameters and known module labels. In this framework, cells were assigned to either a control group or a disease group, and genes were assigned to one of eight predefined rhythm-state modules corresponding to non-rhythmic, stable rhythm, rhythm gain, rhythm loss, amplitude-enhanced rhythmic, amplitude-attenuated rhythmic, phase-advanced rhythmic, and phase-delayed rhythmic patterns.

At the cell level, each simulated cell was assigned a true circadian phase 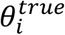 and an observed noisy phase 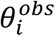. True phases were generated either from a circular uniform distribution or from a von Mises mixture distribution; in the main simulations, true phases were sampled from a von Mises mixture with six phase centers to mimic clustered phase structure. To introduce heterogeneous phase reliability, each cell was assigned a confidence variable

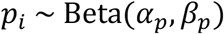

which was mapped to a cell-specific von Mises concentration parameter *k*_*i*_. Observed phases were then generated as

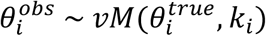

so that cells with higher *p*_*i*_ had lower phase noise. Each cell was additionally assigned a log-normal library-size factor.

At the gene level, each gene *g* was assigned a baseline log-expression *M*_*g*_, a baseline rhythmic amplitude *A*_*g*_, a baseline phase *ϕ*_*g*_, and a negative-binomial dispersion parameter α_*g*_. Group-specific amplitudes and phases were then derived according to the assigned module. For module 0, amplitudes were set to zero in both groups. For module 1, amplitudes and phases were shared across groups. For modules 2 and 3, rhythmicity was present in only one group, corresponding to rhythm gain and rhythm loss, respectively. For modules 4 and 5, amplitudes differed between groups according to predefined log_2 fold-change parameters, while phase differences were constrained to remain small. For modules 6 and 7, amplitudes were preserved across groups whereas phases were shifted by a predefined amount in the negative or positive direction, corresponding to phase advance and phase delay.

Given the true phase and group-specific rhythmic parameters, the expected expression of gene *g* in cell *i* was defined as

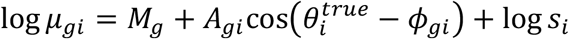

where *A*_*gi*_ and *ϕ*_*gi*_ denote the group-specific amplitude and phase for the group to which cell *i* belongs, and *s*_*i*_ is the cell-specific size factor. Counts were generated from a negative binomial distribution with gene-specific dispersion,

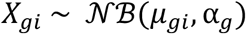

followed by expression-dependent dropout. Specifically, the dropout probability was modeled as a logistic function of the expected expression level, and counts were set to zero accordingly. This simulation framework yields known truth for gene-level baseline expression, amplitude, phase, amplitude change, phase shift, and module label, and was used to evaluate the downstream rhythmic parameter estimation and module-classification components of scRhythmBayes.

### Evaluation framework

Using the simulation frameworks described above, we evaluated scRhythmBayes at three linked levels: cell-level phase inference, gene-level rhythmic modeling and differential rhythm-state classification. Whenever applicable, benchmarking was performed globally and in stratified analyses, including cell type–specific, condition-specific and module-specific settings.

### Evaluation of cell-level phase inference

Using the cell-level phase-inference simulation, we evaluated whether scRhythmBayes accurately recovered continuous cell-level circadian time and whether its posterior confidence summaries were informative for downstream use. Because circadian time is periodic on a 24-h cycle, phase error was quantified using circular absolute error. For a cell with ground-truth time 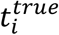 and predicted time 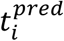, the circular absolute error was defined as

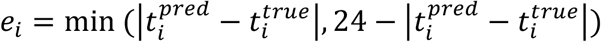

so that errors were constrained to [0,12] h and wrap-around consistency at the cycle boundary was preserved.

To evaluate phase recovery beyond a single-point summary, we used the empirical cumulative distribution function (ECDF) of circular error,

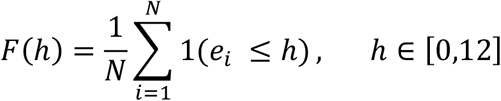

which gives the fraction of cells recovered within an error tolerance h. This representation was used to quantify tolerance-dependent phase accuracy across the full admissible error range rather than at a single threshold. We summarized this curve by its normalized area under the curve,

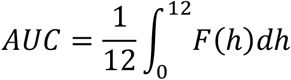

so that AUC is scaled to [0,1], with larger values indicating better phase recovery over a broad range of tolerated errors. Median absolute error (MedAE) was used as the primary summary of typical prediction error, with MAE, RMSE, and circular MAPE reported as complementary summaries. Performance was assessed on the pooled dataset and in cell type–resolved analyses. Posterior resultant length and structured-component responsibility were additionally examined as summaries of phase concentration and confidence.

### Evaluation of gene-level rhythmic modeling

Using the gene-level rhythmic simulation, we evaluated gene-level rhythmic modeling from two complementary perspectives: recovery of simulated rhythmic parameters and identification of rhythmic genes. For parameter recovery, inferred and true values of log-scale baseline, mean expression, amplitude, and phase were compared within each condition and, when appropriate, after pooling across conditions. Linear parameters were assessed via absolute-error metrics, whereas phase was quantified using wrapped circular error. For each parameter, recovery error was summarized using the ECDF of the corresponding error distribution and its normalized AUC. These summaries were supplemented by MedAE, MAE, RMSE, MAPE and *R*^2^.

For rhythmic gene detection, genes were compared against known simulation truth derived from the generating amplitude structure. Truth sets were defined at both the condition-specific and pooled levels, including a detectable-truth setting restricted to genes with sufficiently large simulated amplitude. Detection performance was summarized using sensitivity, specificity, precision, recall, F1 score, and accuracy. To reduce dependence on a single threshold, we additionally performed score-based ROC analysis using method-specific continuous evidence scores and summarized discrimination by ROC-AUC.

### Evaluation of differential rhythm state classification

Using the known eight-module truth from the gene-level simulation, we evaluated whether cross-condition rhythm changes were assigned to the correct predefined biological state. Because these modules represent structured rhythm states rather than unconstrained clusters, evaluation focused on state recovery. Agreement between predicted and true labels was summarized by confusion-matrix analysis, and module-wise one-versus-rest performance was quantified using accuracy, precision, recall and F1 score. Because scRhythmBayes returns posterior probabilities over candidate states, we additionally assessed module discrimination using one-versus-rest ROC analysis based on posterior module probabilities, summarized by ROC-AUC. To separate state-assignment error from parameter-estimation error, parameter recovery was also examined after stratifying genes by true or predicted module membership. When comparison involved methods defined on a reduced state space, such as dryR, simulation truth was collapsed to the closest compatible representation before evaluation.

### Robustness and runtime benchmarks

Robustness analyses were used to test whether model performance and downstream conclusions remained stable under perturbation of key input-design factors and modeling assumptions. For the seed-gene perturbation benchmark, cell-level phase inference was repeated using seed panels containing 27, 20, 15, 10 or 5 seed genes. For the zeitgeber-prior sensitivity benchmark, the default setting using the sample-level zeitgeber prior was compared with a no-prior setting in which the informative zeitgeber anchor was removed or made uninformative. Phase-inference performance in these benchmarks was evaluated using predicted-versus-true circadian time scatter plots, circular absolute error, MedAE and normalized AUC of the circular-error ECDF.

To evaluate temporal-sampling robustness, simulated datasets were analyzed using 24, 12, 8, 6, 4 or 2 input zeitgeber time points across the 24-h cycle. To evaluate cell-number robustness, simulated datasets containing 20,000, 10,500, 6,000, 3,000, 1,500 or 600 cells were analyzed. For both benchmark series, scRhythmBayes was evaluated across cell-phase inference, gene-level rhythmic parameter estimation and differential rhythm-state assignment. Cell-phase recovery was summarized by circular-error ECDFs, normalized AUC and MedAE. Gene-level recovery was summarized for amplitude and phase using *R*^2^, RMSE, MSE, mean error, MAE and MAPE. Module recovery was assessed using confusion matrices comparing predicted and ground-truth rhythm-state modules, together with ground-truth-module-wise proportions of predicted module assignments.

Runtime was measured for the two main computational steps, cell-phase inference and gene-level rhythmic parameter estimation. Runtime was recorded across the same temporal-sampling and cell-number settings and reported in minutes. Module-classification runtime was not included in this benchmark because the primary computational cost was expected to arise from genome-wide gene-level rhythmic model fitting.

For module classification, additional robustness was examined by repeating seed-based initialization over a grid of threshold settings and quantifying assignment consistency, posterior stability and preservation of module-level conclusions. These analyses were used to distinguish genuine model stability from dependence on a single seed set, prior setting, temporal sampling design, cell number or threshold configuration.

### Baseline methods and comparison settings

Baseline comparisons were performed separately for cell-level phase inference and gene-level rhythmic modeling. Cell-level phase benchmarks included Tempo, CYCLOPS and Cyclum^4-6^. Gene-level benchmarks included JTK_CYCLE, harmonic regression and dryR^7-9^. These methods were selected to cover widely used and technically distinct approaches for phase inference and rhythmicity analysis. Each method was run in its native input and output space whenever possible, after which outputs were mapped to a common interpretable scale for evaluation. Phase benchmarking was performed on matched cells across methods, whereas gene-level benchmarking was performed on matched genes using native significance scores, parameter estimates or compatible reduced state representations as appropriate. This design was intended to preserve conceptual comparability while minimizing artifacts introduced by method-specific output formats.

## Data progression and preprocessing

Public single-cell datasets were processed into analysis-ready Seurat objects from raw count matrices^16^. Shared preprocessing steps included cell-level quality control, normalization, dimensionality reduction, clustering, and marker-based cell-type annotation; additional dataset-specific steps, such as doublet removal, batch correction, or multimodal integration, were applied when required by the original data modality and study design. For all downstream scRhythmBayes analyses, raw RNA count matrices were used as input to the modeling framework.

### Liver applications

We analyzed two mouse liver single-cell datasets representing complementary circadian designs^10^. In the primary dataset, cells were sampled at discrete zeitgeber times spanning the 24-h cycle and annotated into periportal and pericentral hepatocyte populations. In the independent validation dataset^12^, hepatocyte populations were used to assess reproducibility of subtype-associated circadian structure.

For the Hep.CV versus Hep.PV comparison, raw hepatocyte counts, numeric zeitgeber labels and hepatocyte subtype annotations were supplied to the phase-inference layer. Cell-level phase estimates and confidence summaries were then propagated to gene-level rhythmic modeling and downstream rhythm-state module analysis. For the multi-cell-type liver analysis, inferred phase summaries and cell-type-resolved gene-level rhythmic parameters were used for SRB_OVR-based rhythmic marker discovery across major liver cell populations.

### Cardiac application

For the heart application, we analyzed a human single-cell dataset from dilated cardiomyopathy containing two clinical rhythm states^17^, normal circadian rhythm (NCR) and abnormal circadian rhythm (ACR), with limited sampling times. The original object was processed through quality control and RNA-based clustering, and major cardiac cell populations were retained when they had sufficient representation across conditions. These populations included cardiomyocytes, fibroblasts, endothelial cells, myofibroblasts, pericytes, smooth muscle cells, myeloid cells, lymphocytes, neurons and adipocytes.

The heart application was designed primarily as a cell-phase-level prioritization analysis in a multi-cell-type, multi-condition disease setting. The relatively low sequencing depth of this dataset was considered when interpreting representative gene trajectories, but the use of the phase-prioritization workflow was motivated by the study design rather than by sequencing depth alone. Phase inference was performed using raw RNA counts, numeric zeitgeber time labels and relevant stratum information. The inferred cell phases were then used to compare temporal organization between NCR and ACR across cardiac cell types.

For each cardiac cell type, inferred phases were summarized in two complementary ways. First, sample-level mean resultant length, ρ, was computed within each biological sample to quantify within-sample phase coordination. Group-wise comparisons of ρ between NCR and ACR were then performed across samples. Second, normalized phase-distribution profiles were constructed for each cell type and group, and DTW-based phase-distribution alignment was used to quantify between-condition phase-profile divergence.

These phase-informed summaries were used to prioritize cardiac cell types showing condition-associated temporal disorganization. Neurons, smooth muscle cells and cardiomyocytes were selected for main-text representative trajectory visualization because they showed informative phase-level patterns and correspond to biologically interpretable axes of cardiac regulation: neuro-cardiac timing, vascular tone remodeling and myocardial contractile-electrophysiological function. Other cardiac cell types were retained in the systematic phase-level analysis and are reported in Supplementary Fig. 11.

## Abbreviations

ZT: zeitgeber time
OVR: one-versus-rest
SRB_OVR: scRhythmBayes one-versus-rest rhythmic marker analysis
DTW: dynamic time warping
AUC: area under the curve
MedAE: median absolute error
RMSE: root mean squared error
MAE: mean absolute error
FDR: false discovery rate
ROC: receiver operating characteristic
GPCR: G protein-coupled receptor
Hep.CV: central-vein hepatocytes
Hep.PV: portal-vein hepatocytes
NCR: normal circadian rhythm
ACR: abnormal circadian rhythm
DCM: dilated cardiomyopathy
BH: Benjamini–Hochberg
ZINB: zero-inflated negative binomial
LRT: likelihood-ratio test
CI: confidence interval
MAPE: mean absolute percentage error
MSE: mean squared error
GAM: generalized additive model
UMAP: uniform manifold approximation and projection.

## Data availability

The public datasets analysed in this study are available from the Gene Expression Omnibus under accession numbers GSE288579 and GSE145197^10,12^. The human dilated cardiomyopathy single-cell dataset analysed in this study was obtained from the published study by Jia et al. The gene universe used for simulation was derived from GSE11923^27^. Reactome pathway annotations used for enrichment analysis are available from the Reactome database. Simulated datasets, processed intermediate files and source data required to reproduce the main figures will be deposited in a public repository upon publication. Additional data supporting the findings of this study are available from the corresponding authors upon reasonable request.

## Code availability

The scRhythmBayes R package and analysis scripts used in this study are available at GitHub: https://github.com/Liwei-DynamicsLab/scRhythmBayes. The repository includes the package source code, example data, documentation and scripts for reproducing the analyses and generating the figures. The version of the package used for this study will be specified in the archived release.

## Acknowledgements

This work was supported by grants from the Natural science foundation of Shandong Province (Grant No. ZR2024MH343). CP.M.-J. was supported by internal funds from the Helmholtz Pioneer Campus, and by the Generalitat Valenciana through the Plan GenT (Project-ID CIDEXG/2023/30). CP.M.-J., I.K.D, L.Z were also supported by the Agencia Estatal de Investigación through the ATRAE program (Project-ID ATR2024-154995), and the Spanish Ministry of Science and Innovation (Project-ID PID2024-157730OB-I00). K.A.D. is supported by Helmholtz Munich, the German Center for Diabetes Research (DZD), and by German Research Foundation (DFG) grant 536691227 within the Research Unit FOR 5795-1 (HyperMet).

## Author contributions

L.Z. conceived the theoretical and mathematical framework of scRhythmBayes and designed the overall methodological strategy. Q.Z. implemented the computational framework, developed analysis workflows, conducted real-data analyses, generated figures, and wrote the article. C.L. contributed to data curation, performed benchmark analyses, validated results, reviewed figures, and contributed to article revision. I.K.D., C.P.M.-J., and K.A.D. provided expert guidance on circadian biology, critically reviewed the analyses and interpretation, and contributed substantial professional feedback on the liver and heart rhythmic programs. W.J., W.L., and Z.W. contributed to data organization, technical support, project discussion, and article revision. L.Z. and P.S. supervised the study, provided project guidance, coordinated the research direction, and contributed to article revision. All authors reviewed and approved the final article.

