## Supplementary Figures for "Decoding Cell-level Circadian Phase and Rhythm-state Remodeling in Single-cell Transcriptomes With scRhythmBayes"

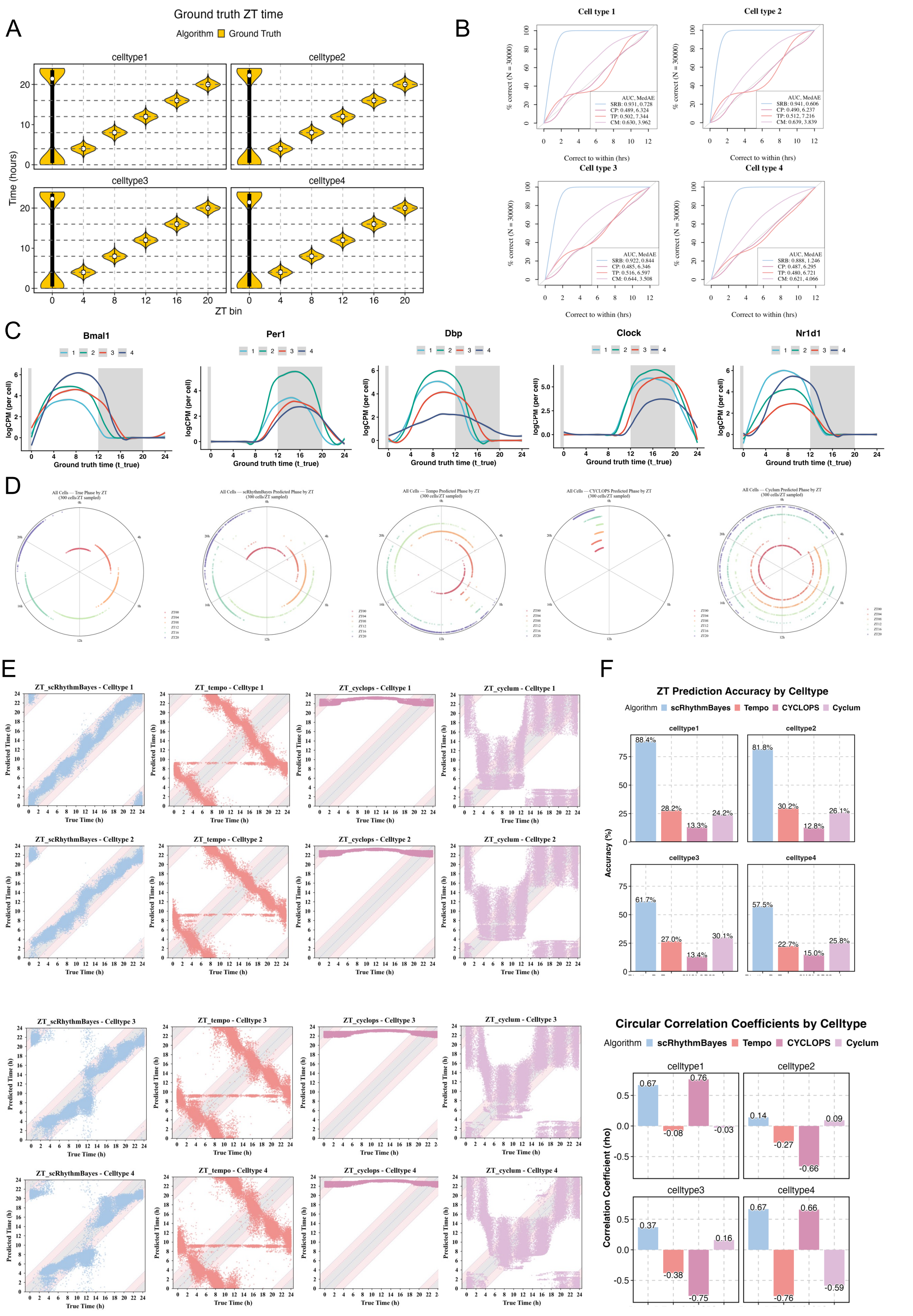


**Supplement. 1 Cell-type-resolved benchmarking of circadian phase prediction in simulated single-cell data.**

**a,** Ground-truth cell-level circadian time distributions across six symbolic sampling bins, shown separately for the four simulated cell types. **b,** Circular absolute error CDFs for each simulated cell type, with normalized AUC and MedAE shown in the insets. **c,** Simulated expression trajectories of representative clock and clock-controlled genes across circadian time in the four simulated cell types. **d,** Circular summaries of phase distributions for Ground Truth and competing methods in each simulated cell type. **e,** Predicted-versus-true circadian time scatter plots for scRhythmBayes, Tempo, CYCLOPS, and Cyclum in each simulated cell type. **f,** Cell-type-specific prediction accuracy and circular correlation coefficients for each method.

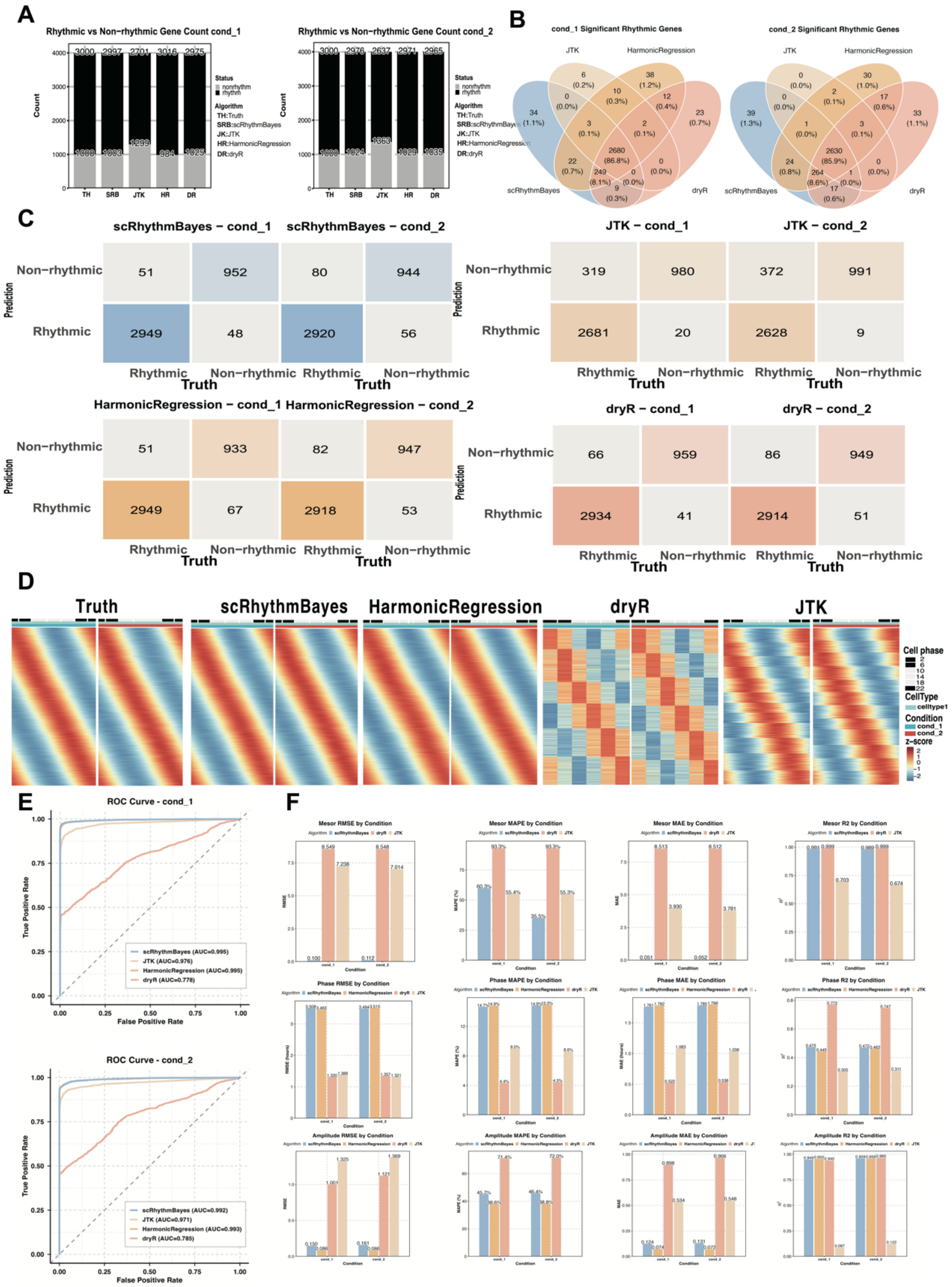


**Supplement. 2 Condition-resolved benchmarking of gene-level rhythmic modeling in simulated single-cell data.**

**a,** Rhythmic and non-rhythmic gene counts in condition1 (cond_1) and condition2 (cond_2) for Ground Truth, scRhythmBayes (SRB), JTK, harmonic regression (HR), and dryR (DR). **b,** Overlap of significant rhythmic genes identified in cond_1 and cond_2 across scRhythmBayes, JTK, harmonic regression and dryR. **c,** Confusion matrices comparing predicted and true rhythmic versus non-rhythmic gene labels in cond_1 and cond_2 for scRhythmBayes, JTK, harmonic regression and dryR. **d,** Representative heatmaps of rhythmic gene patterns for Truth, scRhythmBayes, harmonic regression, dryR and JTK in celltype1 across the two simulated conditions. **e,** ROC curves for rhythmic gene detection in cond_1 and cond_2. **f,** Condition-wise performance summaries for mesor, phase and amplitude recovery, including MAE, MAPE, RMSE and $R^2$ for each method in cond_1 and cond_2.

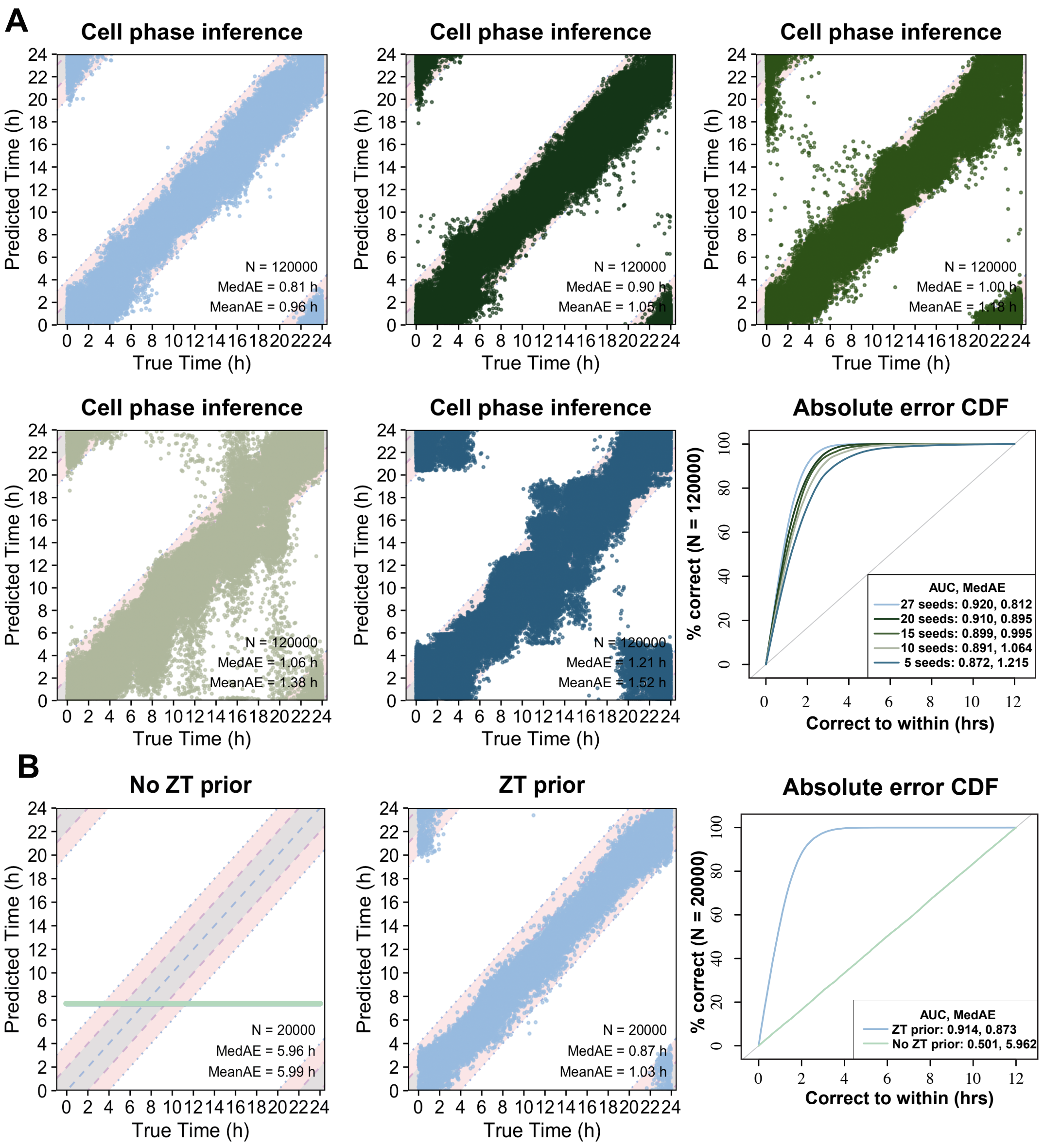


**Supplement. 3 Seed-gene perturbation and zeitgeber-prior sensitivity analyses for cell-level circadian phase inference.**

**a,** Seed-gene perturbation benchmark for scRhythmBayes phase inference using 5, 10, 15, 20 and 27 seed genes. Predicted versus true circadian time and circular absolute error cumulative distribution functions are shown for each seed-gene setting, with normalized area under the curve (AUC), median absolute error (MedAE), mean absolute error (MeanAE) and cell number indicated. **b,** Zeitgeber-prior sensitivity benchmark comparing scRhythmBayes phase inference with and without the sample-level zeitgeber time prior. Predicted versus true circadian time and circular absolute error cumulative distribution functions are shown for the two settings, with AUC, MedAE, MeanAE and cell number indicated.

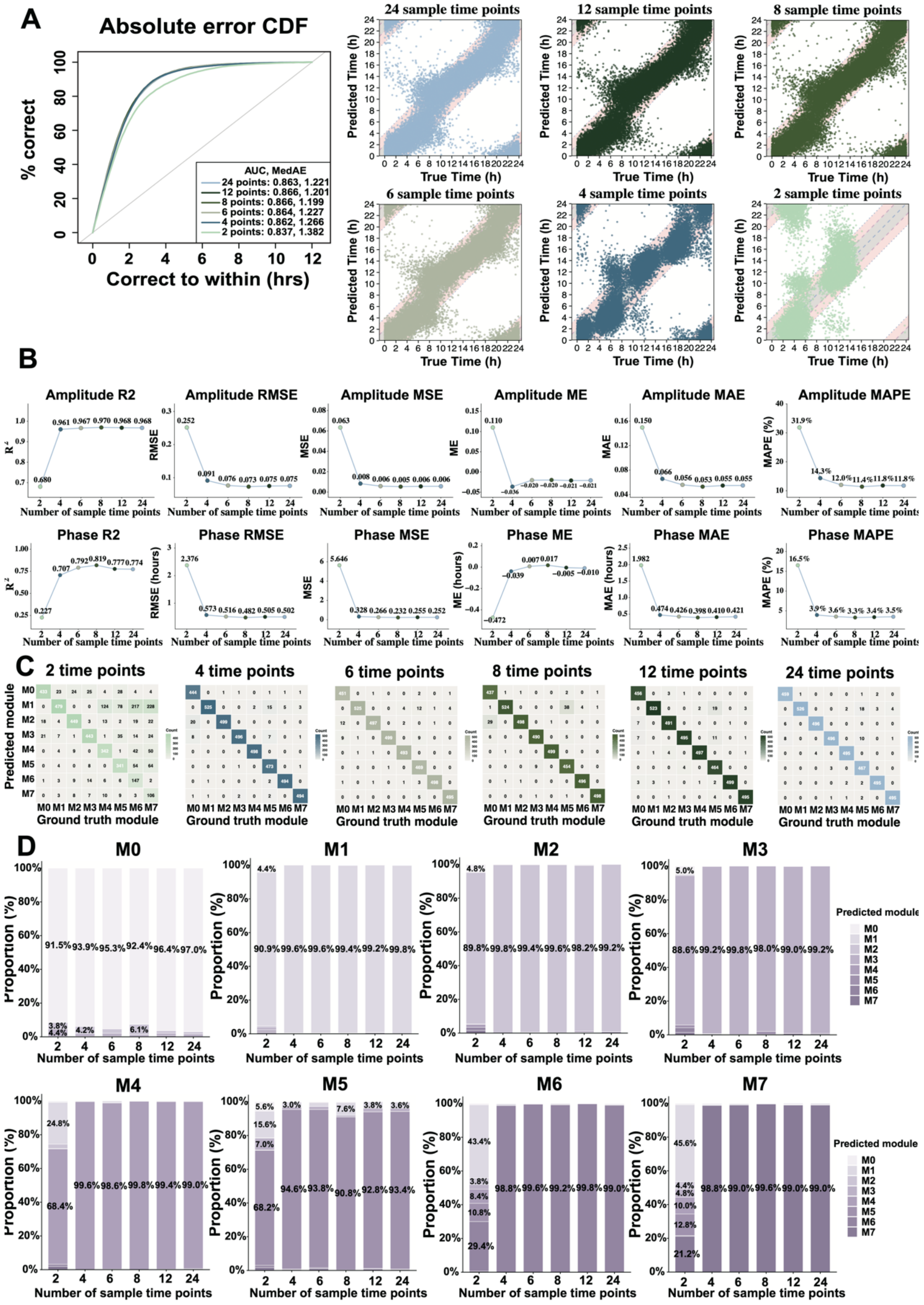


**Supplement. 4 Robustness of scRhythmBayes to reduced input zeitgeber time points in simulated single-cell data.**

**a,** Cell-phase inference across 2, 4, 6, 8, 12 and 24 input zeitgeber time points, shown by circular absolute error cumulative distribution functions and predicted versus true circadian time. Insets report normalized area under the curve (AUC) and median absolute error (MedAE). **b,** Gene-level amplitude and phase recovery across sampling-density settings, summarized by $R^2$, root mean squared error (RMSE), mean squared error (MSE), mean error (ME), mean absolute error (MAE) and mean absolute percentage error (MAPE). **c,** Confusion matrices comparing predicted and ground-truth rhythm-state modules across sampling-density settings. **d,** Ground-truth-module-wise proportions of predicted module assignments across sampling-density settings.

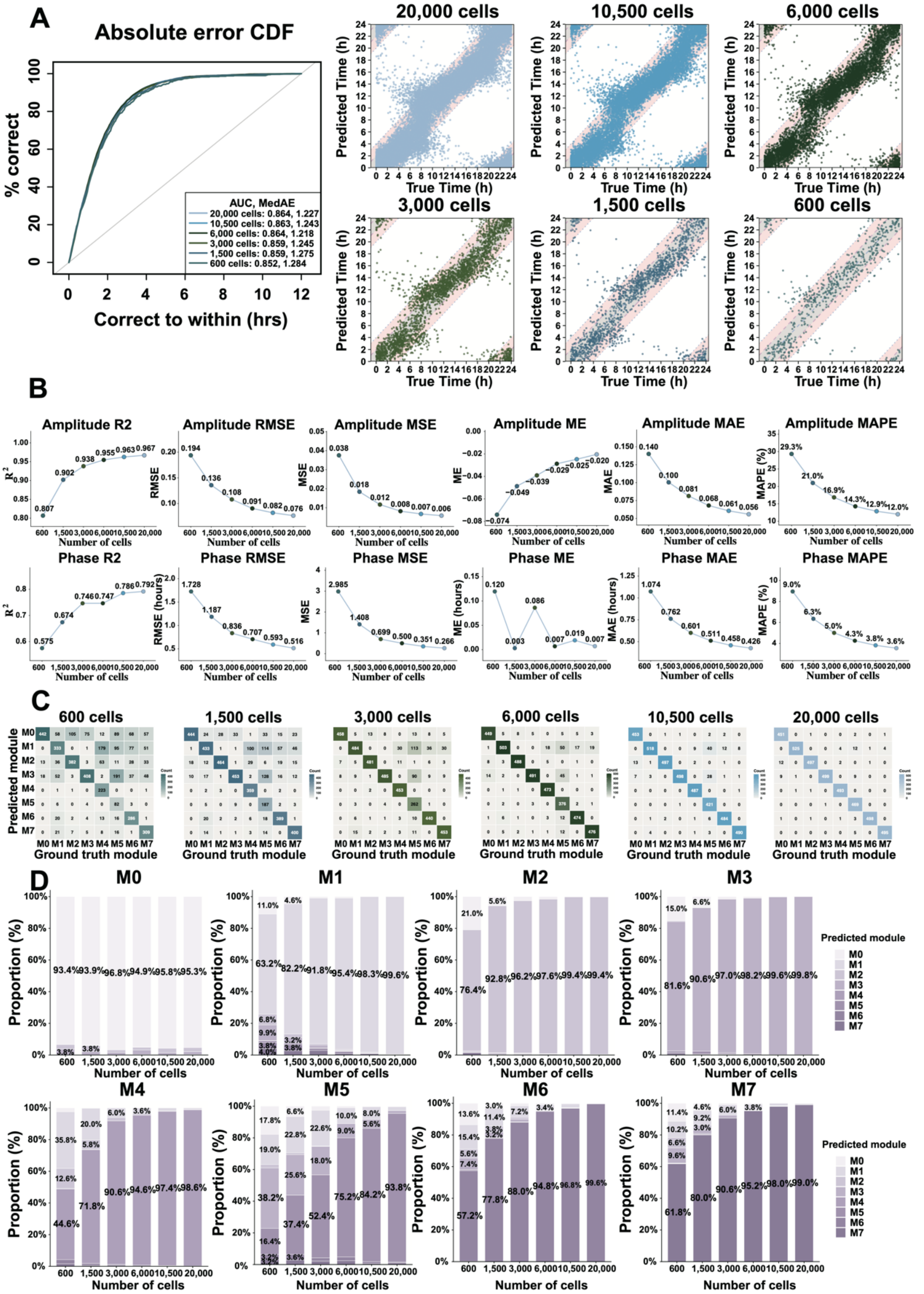


**Supplement. 5 Robustness of scRhythmBayes to reduced cell numbers in simulated single-cell data.**

**a,** Cell-phase inference across simulated datasets containing 20,000, 10,500, 6,000, 3,000, 1,500 and 600 cells, shown by circular absolute error cumulative distribution functions and predicted versus true circadian time. Insets report normalized area under the curve (AUC) and median absolute error (MedAE).

**b,** Gene-level amplitude and phase recovery across cell-number settings, summarized by $R^2$, root mean squared error (RMSE), mean squared error (MSE), mean error (ME), mean absolute error (MAE) and mean absolute percentage error (MAPE). **c,** Confusion matrices comparing predicted and ground-truth rhythm-state modules across cell-number settings. **d,** Ground-truth-module-wise proportions of predicted module assignments across cell-number settings.

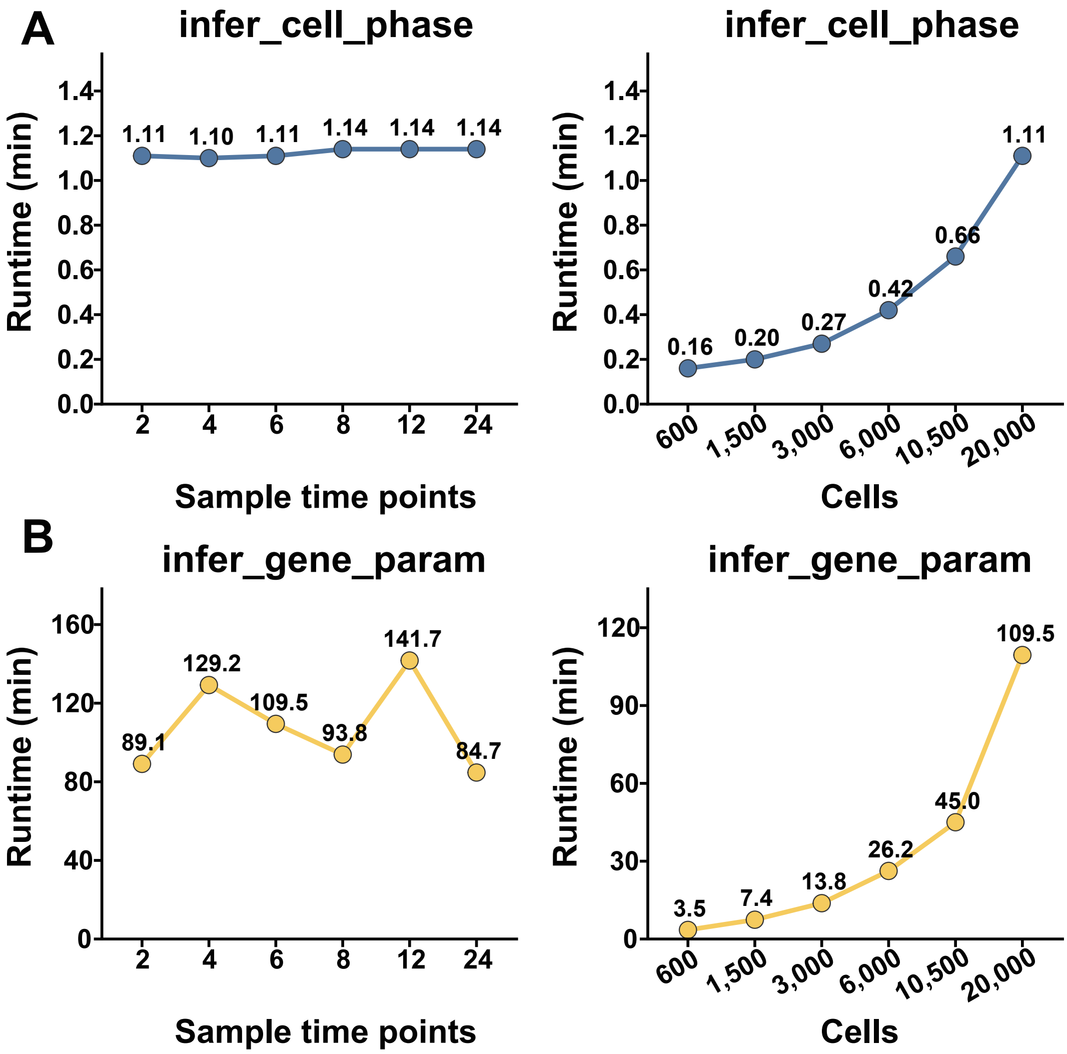


**Supplement. 6 Runtime of scRhythmBayes phase inference and gene-level rhythmic parameter estimation.**

**a,** Runtime of infer_cell_phase across 2, 4, 6, 8, 12 and 24 input sample time points and across 600, 1,500, 3,000, 6,000, 10,500 and 20,000 cells. **b,** Runtime of infer_gene_param across the same sampling-density and cell-number settings. Runtime is reported in minutes.

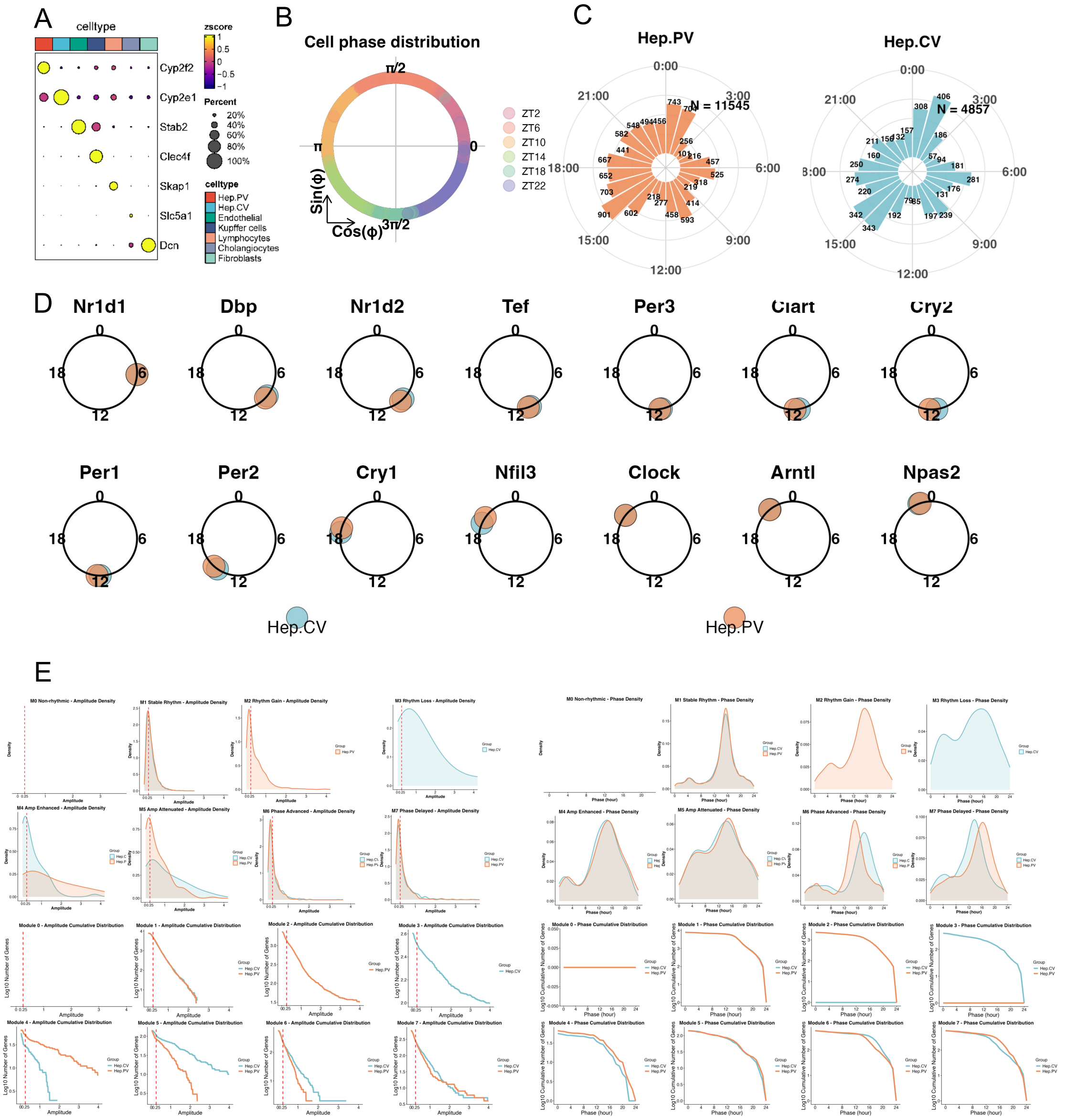


**Supplement. 7 Phase structure and module-level rhythmic features in two hepatocyte subtypes.**

**a,** Marker-gene dot plot showing the annotated liver cell populations, including the two hepatocyte subtypes analyzed downstream. **b,** Circular summary of predicted phase occupancy across rounded ZT groups. **c,** Phase distributions in Hep.PV and Hep.CV shown on a 24-h circular scale. **d,** Circular summaries of representative core clock genes in the two hepatocyte subtypes. **e,** Module-wise density distributions and cumulative distributions of amplitude and phase features in Hep.CV and Hep.PV.

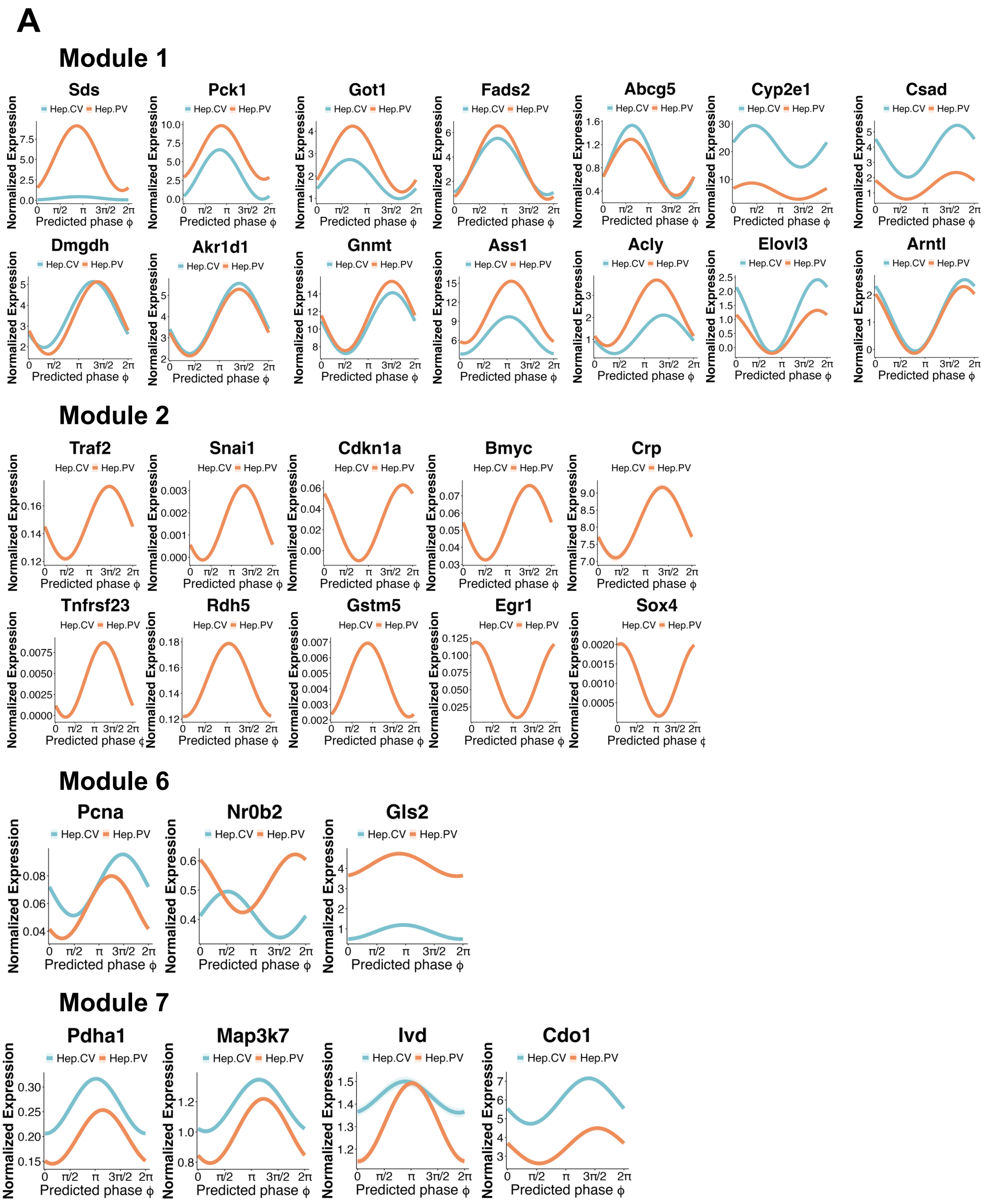


**Supplement. 8 Representative expression trajectories of module-associated genes underlying the drug-metabolism-related programs in two hepatocyte subtypes.**

**a,** Representative phase-ordered expression trajectories of genes from Module 1, Module 2, Module 6 and Module 7 across predicted phase in Hep.CV and Hep.PV.

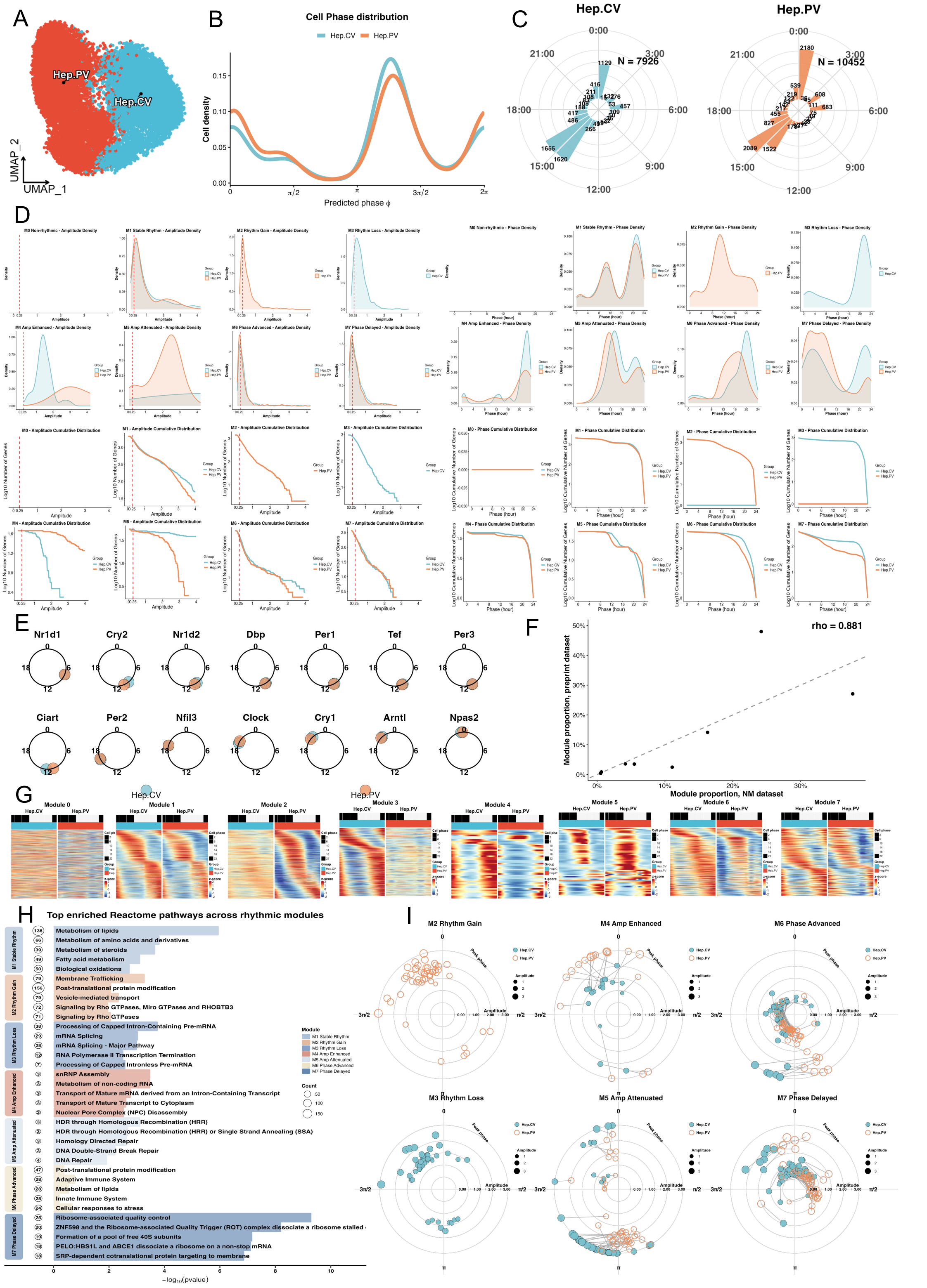


**Supplement. 9 Application of scRhythmBayes to circadian programs in two hepatocyte subtypes in an independent hepatocyte single-cell dataset.**

**a,** UMAP of the independent hepatocyte single-cell dataset (GSE145197), showing Hep.CV and Hep.PV hepatocytes. **b,** Density distributions of predicted phase in Hep.CV and Hep.PV. **c,** Circular phase distributions in Hep.CV and Hep.PV on the 24-h scale. **d,** Module-wise density distributions and cumulative distributions of amplitude and phase features in Hep.CV and Hep.PV. **e,** Circular summaries of representative core clock genes in the two hepatocyte subtypes. **f,** Cross-dataset comparison of rhythm-state module proportions between the primary and independent hepatocyte datasets. **g,** Paired heatmaps of gene expression patterns across the eight scRhythmBayes modules in the two hepatocyte subtypes. **h,** Top enriched Reactome pathways across rhythmic modules. **i,** Polar plots of representative genes from differential rhythm modules, including rhythm gain, rhythm loss, amplitude-enhanced, amplitude-attenuated, phase-advanced and phase-delayed patterns in Hep.CV and Hep.PV.

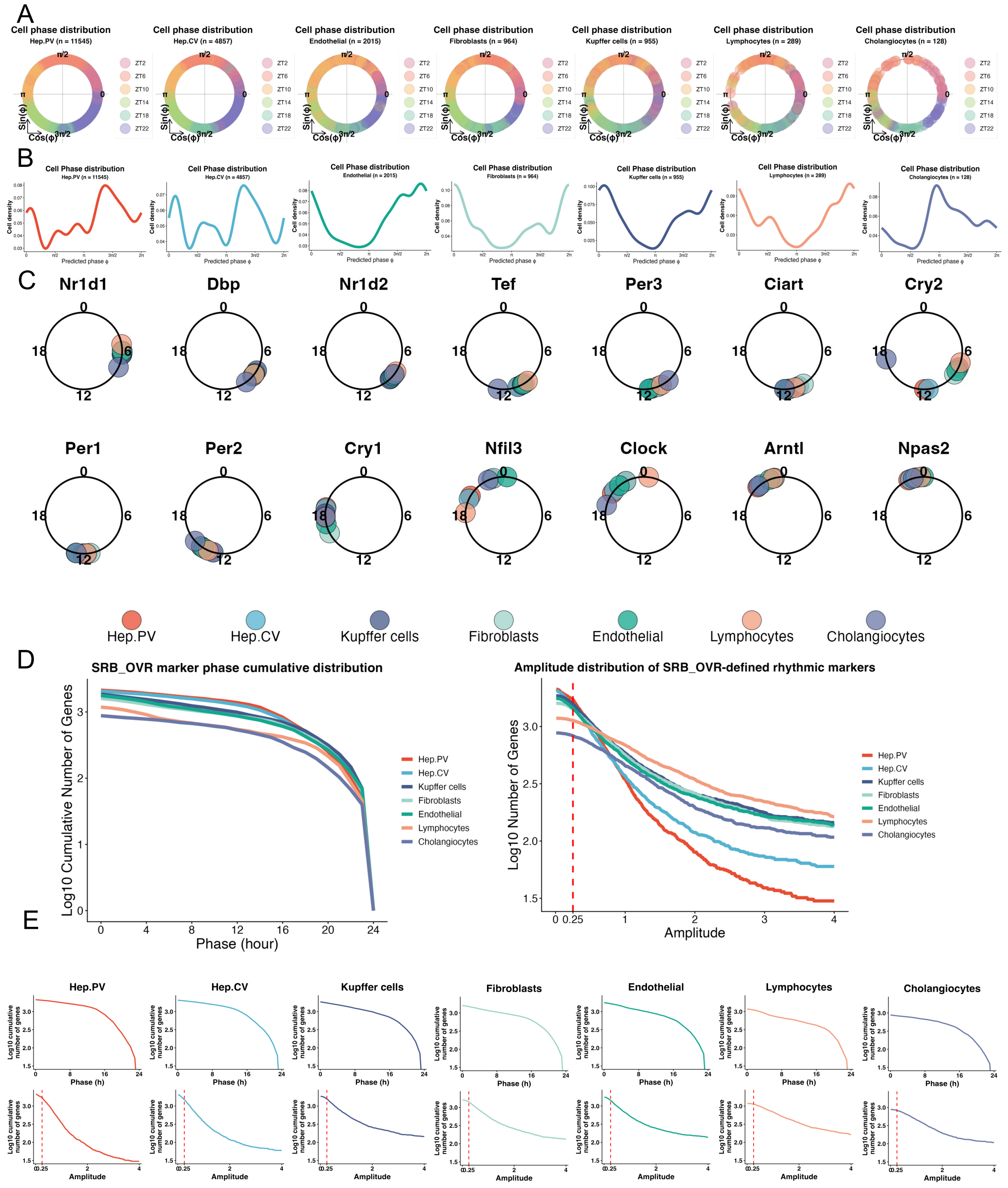

**Supplement. 10 Cell-type-resolved phase organization and SRB_OVR-defined rhythmic marker features across liver cell types.**

**a,** Circular summaries of scRhythmBayes-inferred cell phase distributions across major liver cell types, including Hep.PV, Hep.CV, endothelial cells, fibroblasts, Kupffer cells, lymphocytes and cholangiocytes. Colors indicate rounded Zeitgeber time groups. **b,** Density distributions of predicted cell phase for each liver cell type, showing cell-type-specific phase occupancy across the circadian cycle. **c,** Circular phase summaries of representative core clock genes across liver cell types. Each point represents the estimated phase of a clock gene in one liver cell type, with colors indicating cell-type identity. **d,** Combined cumulative phase distribution and amplitude distribution of SRB_OVR-defined cell-type-specific rhythmic markers across liver cell types. The left panel shows cumulative marker-gene counts across predicted phase on the 24-h scale, and the right panel shows cumulative marker-gene counts across rhythmic amplitude. The red dashed line indicates the minimum-amplitude threshold used for marker filtering. **e,** Cell-type-resolved cumulative phase and amplitude distributions of SRB_OVR-defined rhythmic markers. For each cell type, the upper panel shows the cumulative phase distribution and the lower panel shows the corresponding amplitude distribution, enabling comparison of phase coverage and rhythmic amplitude structure across cell-type-specific rhythmic programs.

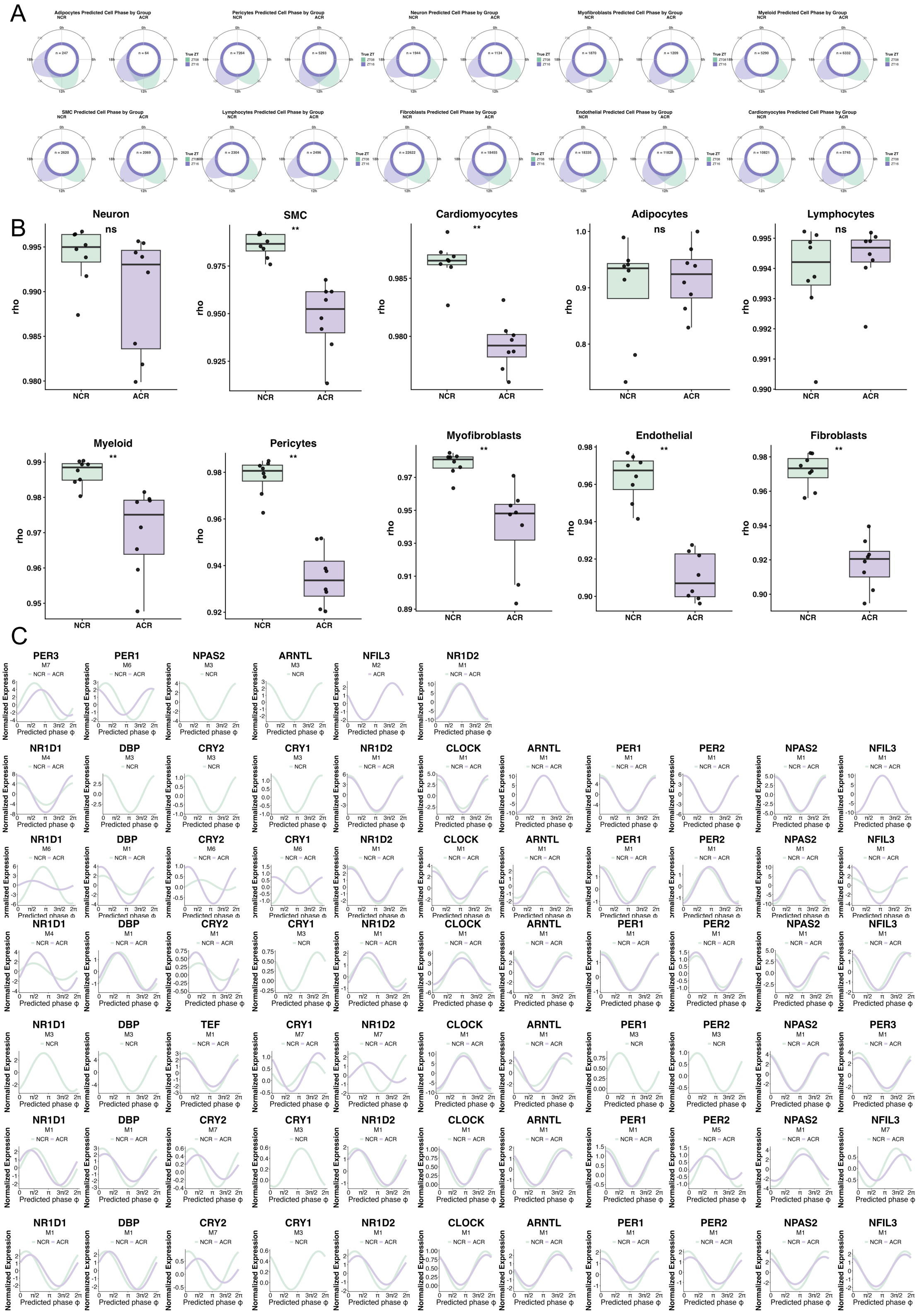


### Supplement. 11 Cell-type-resolved phase distributions, sample-level phase coordination and representative gene trajectories across major cardiac cell types.

**a,** Circular summaries of scRhythmBayes-inferred cell phase distributions in normal circadian rhythm (NCR) and abnormal circadian rhythm (ACR) groups across major cardiac cell types. Each circular plot shows one cell type within one condition on the 24-h phase scale. **b,** Cell-type-wise comparison of sample-level mean resultant length, $\rho$, between NCR and ACR. Each point represents one biological sample within a given cell type; higher $\rho$ indicates stronger within-sample phase coordination. Statistical labels indicate group-wise comparisons. **c,** Phase-ordered expression trajectories of representative core clock and cell-type-relevant genes across additional cardiac cell types. Rows are arranged, from top to bottom, as adipocytes, lymphocytes, myeloid cells, pericytes, myofibroblasts, endothelial cells and fibroblasts. Curves show normalized expression as a function of scRhythmBayes-inferred phase in NCR and ACR.
